# Repurposing statins and phenothiazines to treat chemoresistant neuroblastoma

**DOI:** 10.1101/2025.01.28.635193

**Authors:** Katarzyna Radke, Kristina Aaltonen, Erick A. Muciño-Olmos, Javanshir Esfandyari, Adriana Mañas, Alexandra Seger, Aleksandra Adamska, Karin Hansson, Oksana Rogova, Daniel J. Mason, Daniel J. O’Donovan, Ian Roberts, Antonia Lock, Jane Brennan, Emma J. Davies, Peter Spégel, Oscar C. Bedoya-Reina, David Brown, Neil T. Thompson, Cesare Spadoni, Daniel Bexell

**Affiliations:** Translational Cancer Research, Lund University; Lund, Sweden; Translational Research in Pediatric Oncology, Hematopoietic Transplantation and Cell Therapy, IdiPAZ Research Center, University Hospital La Paz; Madrid, Spain; Pediatric Onco-hematology Clinical Unit IdiPAZ-CNIO, National Cancer Research Center (CNIO); Madrid, Spain; Cancer Stem Cell Laboratory, The Breast Cancer Now Toby Robins Research Centre, The Institute of Cancer Research; London, UK; Department of Chemistry, Centre for Analysis and Synthesis, Lund University; Lund, Sweden; Healx Ltd.; Charter House, 66-68 Hills Road, Cambridge, CB2 1LA, UK; School of Medical Sciences, Örebro University; 70182 Örebro, Sweden; Childhood Cancer Research Unit, Department of Women’s and Children’s Health, Karolinska Institutet; 11883 Stockholm, Sweden; aPODD Foundation; Virginia Cottage, Bridgnorth Road, Stourton, Stourbridge, West Midlands, DY7 5BQ, UK

**Keywords:** Cancer, chemoresistance, drug repurposing, machine learning, neuroblastoma, phenothiazine, statin, in silico predictions, prochlorperazine

## Abstract

Relapse and treatment resistance are common in children with high-risk neuroblastoma, and novel therapies are needed. Conventional drug discovery is slow, expensive, often fails in practice, and consequently falls short in addressing pediatric and rare conditions. In such instances, drug repurposing is a promising strategy. Here, we used two independent *in silico* prediction tools including machine learning to identify approved drugs for repurposing against neuroblastoma. The combination of statins and phenothiazines showed strong synergistic effects in human neuroblastoma organoids, decreased tumor growth and prolonged survival in *MYCN*-amplified neuroblastoma patient-derived xenografts. The drug combination altered cholesterol metabolism through a dual-hit mechanism and induced a phenotypic switch towards an adrenergic cell state accompanied by increased sensitivity to chemotherapy. Integration of the drug combination into standard-of-care chemotherapy regressed tumors and prolonged survival in chemoresistant patient-derived xenografts. Thus, a combination of safe and approved medications added to standard-of-care chemotherapy outperforms chemotherapy alone in chemoresistant neuroblastoma.

**Teaser:** Using *in silico* prediction tools we repurposed statins and phenothiazines as a combination with translational potential in high-risk neuroblastoma.

## INTRODUCTION

The rarity of pediatric cancer and ethical considerations specific to children have hindered drug development for this group of patients(*1, 2*). One option to progress therapies for rare diseases is drug repurposing, where an approved drug with human safety and efficacy data is given for an alternative, -unapproved indication, which is a timesaving and cost-effective approach. Patient groups with limited treatment options can especially benefit from this approach. Furthermore, efficacy and safety data on existing drugs can provide the basis for computational predictions and help with mechanistic rationalization in screening approaches and preclinical and clinical trials(*3, 4*).

Neuroblastoma (NB) is a childhood malignancy of the sympathetic nervous system responsible for nearly 15% of pediatric cancer-related deaths. NB is a clinically heterogeneous disease, spontaneously regressing with minimal intervention in some children but pursuing an aggressive metastatic course in others. Patients with high-risk NB undergo intensive chemotherapy, surgery, radiotherapy, and other treatments, and, although initial treatment can be effective, over 50% of patients with high-risk NB eventually relapse with treatment-resistant disease. Thus, there is an unmet need for new and better treatment approaches for children with high-risk NB(*5–8*).

The molecular pathology of NB is characterized by copy number aberrations (CNAs) including 1p loss, *MYCN* amplification, 11q loss, 17q gain, and others. Relapsed tumors are enriched for mutations in genes activating oncogenic signaling pathways, such as the RAS–mitogen-activated protein kinase (MAPK) and the YAP-Hippo pathways(*9, 10*). However, compared with many adult tumors, NB has a relatively low mutational burden and few recurrent mutations. While genetic changes, especially CNAs, are important for NB initiation and progression, it has become increasingly clear that transcriptional activity and phenotypic plasticity are important as the disease progresses and fails to respond to treatment(*11*).

Epigenetic and transcriptional analyses have defined two NB cell states *in vitro*; a lineage-committed adrenergic (ADR or ADRN) state, and an immature neural crest cell-like or mesenchymal-like (MES) state(*12, 13*). These cell states exist along a continuum with a high degree of plasticity. Current data suggest that both cell states are important for NB progression, with the MES-like state involved in chemoresistance and relapse(*12–22*).

Disease-gene expression matching (DGEM) represents a methodology aimed at identifying potential therapeutic drugs for diseases characterized by specific gene expression signatures(*23*). By comparing genes differentially expressed in disease states with the gene expression profiles induced by various drugs, existing pharmaceuticals suitable for repurposing can be identified. This approach holds particular relevance for conditions such as NB, where transcriptomic alterations play a critical role in disease progression(*24, 25*).

Another prediction algorithm, PRISM (Predicted Repurposed Indications from Similarity Matrices), can predict novel treatment indications based on known treatment indications and similarity measures between compounds and diseases using machine learning. These measures are obtained by comparing the compounds’ structures or examining their target similarities. Additionally, PRISM defines the similarity between diseases by analyzing ontological data, disease targets, and the frequencies with which diseases co-occur in literature(*26*). Here, PRISM was used to find new drug-disease associations for NB.

We harnessed the power of DGEM and PRISM prediction tools to identify potential candidates for drug repurposing in high-risk NB. This yielded several drug hits already approved for non-cancer diseases. The efficacies of candidate drugs were validated in *MYCN*-amplified chemoresistant human NB organoids and in *MYCN*-amplified patient-derived xenograft (PDX) models. A combination of selected classes of drugs induced significant alterations in cholesterol metabolism and shifted NB cell state signatures towards a chemosensitive ADR-like state *in vitro* with transcriptional features of low-risk NB patient tumors. Notably, treatment with the novel drug combination alongside standard-of-care (COJEC) chemotherapy resulted in tumor regression and extended survival compared with standard-of-care chemotherapy alone in a chemoresistant NB PDX model, paving the way for testing of this combination towards clinical practice.

## RESULTS

We identified drugs for repurposing in high-risk NB using *in silico* prediction tools including machine learning, validation of single drugs and drug combinations in human NB organoids, transcriptional analysis of treatment-induced phenotype, lipidomics, and *in vivo* testing in NB chemoresistant PDX models (**Fig. 1**).

**Fig. 1.**
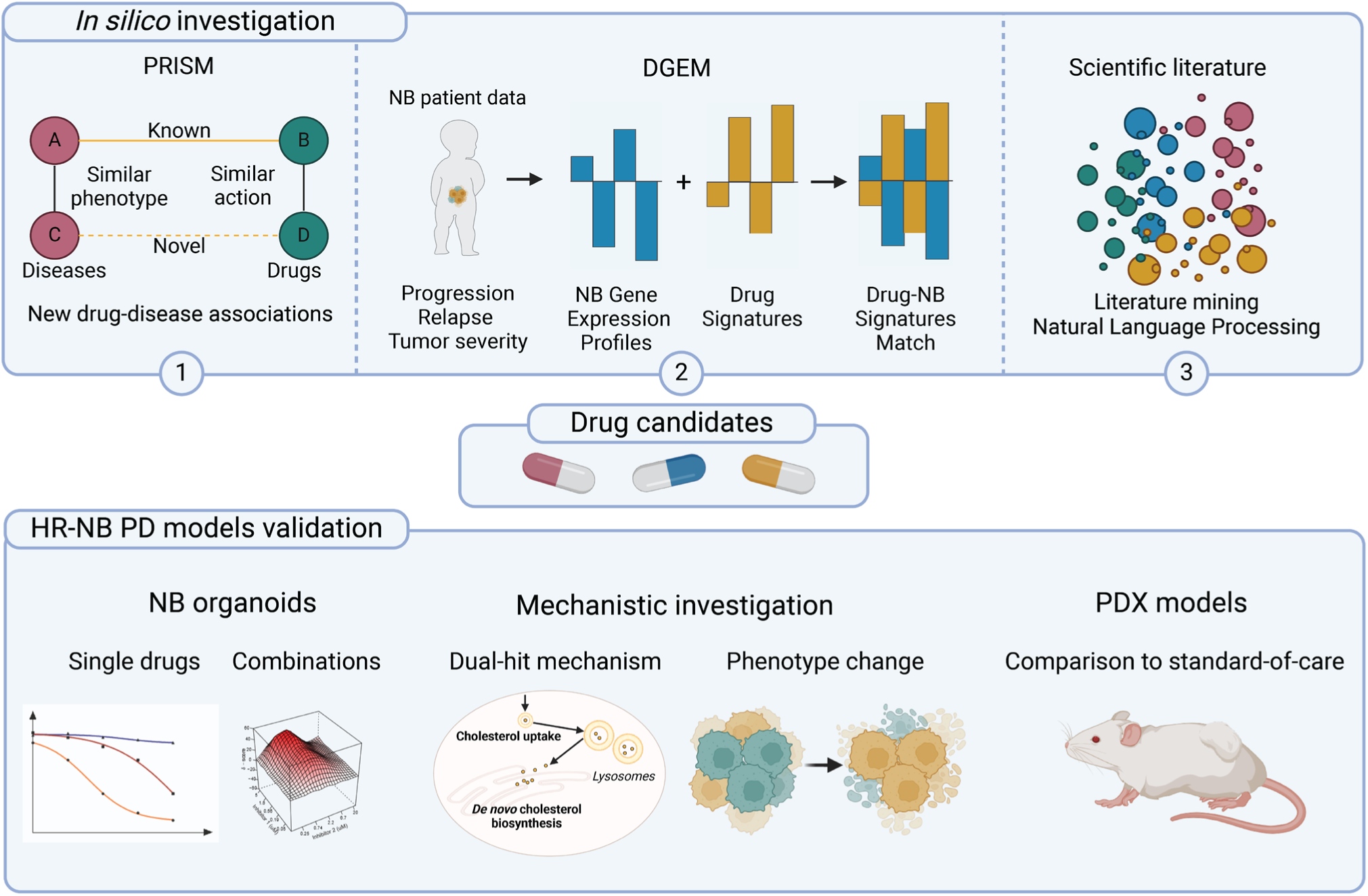
Overview of the study. *In silico*, potential drugs for repurposing were identified using three complementary methods: 1) New drug-disease associations were found based on the PRISM (guilt-by-association) machine learning algorithm, 2) Drug-NB signature matches were identified with DGEM using neuroblastoma gene expression profiles of progression, tumor severity, or relapse, together with a repository of drug signatures, and 3) Literature mining with natural language processing was used as a confirmation tool for hits commonly identified with the other methods. Drug candidates were subsequently tested in high-risk neuroblastoma (HR-NB) models as single drugs and in combination. Investigation of the mechanism of action of the selected drugs identified a dual-hit mechanism and gene expression profile-based phenotype changes after combination treatment. The effect of the combination was investigated *in vivo* along with standard-of-care chemotherapy. HR-NB: high-risk neuroblastoma; DGEM: Disease-Gene Expression Matching; PRISM: Prediction of Repurposed Indications with Similarity Matrices; PD models: patient-derived models.

### In silico drug selection using DGEM and PRISM

Publicly available raw gene expression datasets were manually curated and analyzed for their suitability to reflect the transcriptomic profile of the disease. Four raw gene expression datasets passed quality control criteria that ensured robustness of the input data. These datasets included transcriptomic data from NB patients with associated clinicopathological information including the disease stage, relapse status, and molecular determinants of risk such as *MYCN* amplification and 17q gain (**Table S1, Supplementary Data 1**). Gene expression profiles (GEPs) generated from differentially expressed genes in these datasets were evaluated by gene set enrichment analysis (GSEA) to ensure that they described biological processes indicative of an NB phenotype. All datasets had consistently upregulated terms in keeping with NB biology and clinical data of the studies used to define GEPs such as MYC targets, E2F targets, glycolysis, oxidative phosphorylation, mTORC1 signaling, and proliferation (G2M checkpoint) (**Fig. S1A**). The NB GEPs obtained through this process were used for drug matching.

DGEM predicts drugs likely to be effective in a given disease based on the most differentially expressed genes in a GEP. Briefly, a disease GEP is compared to a compendium of thousands of public and private drug GEP to detect drug-disease associations. The dataset covers a range of pharmacological classes and includes a mixture of approved, experimental, and investigational compounds to guide repurposing recommendations. The strongest drug signatures may, for example, be inversely correlated with the disease signature, in which case the drug may elicit a therapeutic effect on the disease.

In an independent analysis, potential NB targeting drugs were also identified with a guilt-by-association approach using the PRISM machine learning algorithm. Here, compound similarity measures like structural similarities and target homologies, as well as disease similarities retrieved from ontological data, target profiles, and literature co-occurrence measures were used for finding previously unexplored treatment candidates(*26, 27*).

The combined prioritized treatment predictions from DGEM and PRISM were subsequently reviewed by pharmacologists and bioinformaticians, and drug selection was performed based on studies of drug classes, known interactions with pathway, phenotypes, cellular processes, mechanisms, and novel biology. Strength of rationale for efficacy in NB and particular qualities of the drug compounds, including reformulation potential and known safety profiles were accounted for.

Known cancer agents were ranked *in silico* for their likely efficacy against NB, but priority for subsequent laboratory testing was given to safe, non-cancer drugs that might avoid the addition of more toxic chemotherapy agents in NB patients. Twelve substances (**Table 1**) with potential efficacy in NB were selected for further *in vitro* testing.

**Table 1.** Predicted drugs tested in neuroblastoma organoids.

| Drug | Drugbank ID | MW (g/mol) | Known mechanism |
| --- | --- | --- | --- |
| Trifluoperazine (TFP) | DB00831 | 480.42 | **Antipsychotic** drug used in schizophrenia<br>Anti-adrenergic, antidopaminergic<br>**Phenothiazine** |
| Thioridazine (TRD) | DB00679 | 407.04 | **Antipsychotic** drug used in schizophrenia<br>Lower potency than trifluoperazine and withdrawn<br>**Phenothiazine** |
| Nortriptyline | DB00540 | 299.84 | Antidepressant |
| Prochlorperazine (PCZ) | DB00433 | 606.09 | **Antipsychotic** drug,<br>D2 receptor antagonist<br>Anti-emetic properties, **Phenothiazine** |
| Resveratrol | DB02709 | 228.25 | Stilbenoid, natural phenol<br>Dietary supplement |
| Sildenafil | DB00203 | 666.7 | Vasodilation used in e.g., Viagra |
| Dasatinib | DB01254 | 488.01 | Chemotherapy used in e.g. AML and ALL |
| Pazopanib | DB06589 | 437.52 | RTK inhibitor, angiogenesis inhibition<br>Used in e.g., renal cell carcinoma |
| Lovastatin (LOV) | DB00227 | 404.54 | Cholesterol-lowering **statin**<br>Inhibitor of HMG-CoA reductase |
| Folic Acid | DB00158 | 441.4 | Vitamin B |
| Topiramate | DB00273 | 339.36 | Anti-convulsant |
| Tianeptine | DB09289 | 436.95 | Anti-depressant |

### Testing of single drugs in NB organoids

Selected agents were tested as single drugs in three human NB organoid models (LU-NB-1, LU-NB-2, and LU-NB-3) derived from *MYCN*-amplified high-risk PDX tumors (**Table S2**)(*28*). These PDX models retain the genotype, transcriptional profiles, and phenotypes of their corresponding patient tumors(*29*). Four of the 12 drugs decreased the viability of all three NB organoid models (**Fig. 2A; Fig. S1B-D**): the three phenothiazines (used as antipsychotics, dopamine receptor antagonists) trifluoperazine (TFP), thioridazine (TRD), and prochlorperazine (PCZ), and the HMG-CoA reductase inhibitor lovastatin (LOV) (**Fig. 2A**). The half-maximal inhibitory concentration (IC_50_) and area under the curve (AUC) were calculated for these four drugs at 3 and 7 days (**Table S3**). The effect of lovastatin was higher on day 7 compared with day 3 (**Table S3**).

**Fig. 2.**
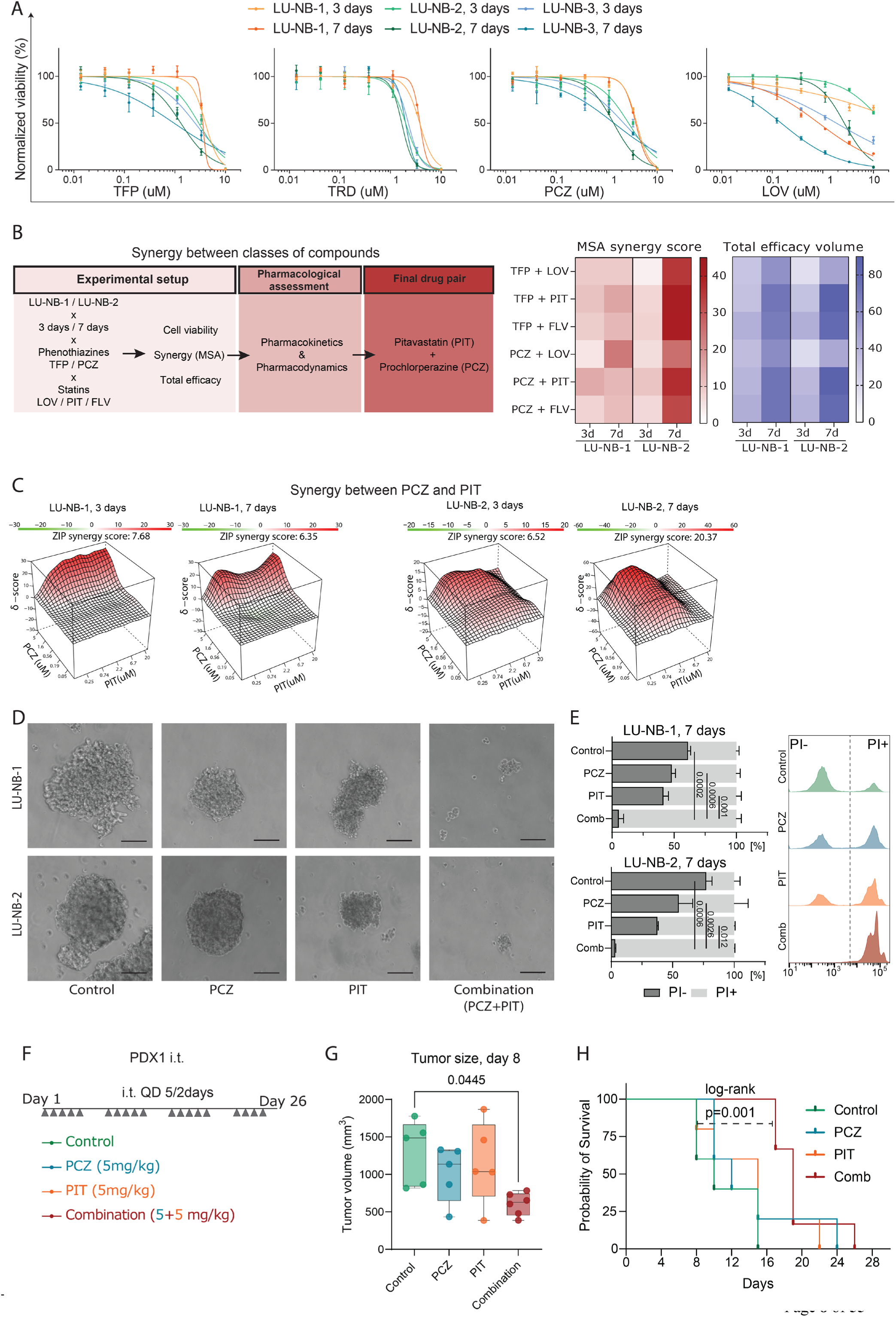
Single drug efficacy and PCZ+PIT combination synergy. **A** Dose-response curves of the four top hits tested in three PDX-derived high-risk neuroblastoma organoids (LU-NB-1, 2 and 3) for 3 and 7 days. **B** Synergy testing between the two drug classes, phenothiazines and statins. Work flow for identifying the final drug pair. Combination total scores of 6 x 6 synergy matrices over 3 or 7 days. **C** Synergy landscape of the PCZ + PIT combination tested in LU-NB-1 and LU-NB-2 organoids for 3 and 7 days. **D** Photographs of NB organoid size and integrity following single drug and PCZ+PIT combination treatment (scale bar 100 µm), 7 days. **E** Propidium iodide (PI) staining for PCZ+PIT combination over a 7-day treatment, (one-way ANOVA, Dunnett’s multiple test). **F** Experimental set-up for intratumoral (i.t.) treatment with single drugs and the combination. Subcutaneous LU-NB-1 PDX tumors were established in NSG mice. **G** Average tumor volume per group, day 8 (n=5 in control, PCZ, and PIT groups; N=7 in combination group, one-way ANOVA followed by Tukey’s multiple comparisons test). **H** Kaplan-Meier curves showing survival in days from the first treatment day. Statistical analysis was performed using the log-rank test between control and combination group. TFP-trifluoperazine; TRD-thioridazine; PCZ-prochlorperazine; LOV-lovastatin; PIT-pitavastatin; FLV-fluvastatin; ZIP-zero interaction potency score.

### Drug combinations are synergistic in NB organoids

Combination therapy with multiple drugs can reduce the development of resistance and reduce the risk of drug toxicity as lower concentrations of individual drug are required(*30*). We hypothesized that combining the two most effective single agent drugs with different mechanisms of action, cholesterol-lowering statins (LOV) and one of the phenothiazines (TFP, TRD, or PCZ), could be a promising combination approach.

We therefore tested combinations of LOV with either TFP or PCZ in the LU-NB-1 and LU-NB-2 organoid models, due to their chemoresistant characteristics *in vitro* and *in vivo*(*19*). Combination treatment in LU-NB-1 and LU-NB-2, respectively, resulted in synergistic ZIP scores for LOV + PCZ (2.84/3.24 at 3 days and 14.3/11.67 at 7 days) and for LOV + TFP (6.36/1.29 at 3 days and 5.03/17.34 at 7 days) (**Fig. S2A**).

To confirm that synergistic effects were not specific to these drugs but more general to the two drug classes, we evaluated the most synergistic area (MSA) and efficacy scores between phenothiazines (TFP and PCZ) and different statins (lovastatin (LOV), fluvastatin (FLV), and pitavastatin (PIT)). To select optimal drugs for further testing and translational relevance, we also examined the *in vitro* combinatory effects and pharmacokinetic features of the drugs together with their intended use. PIT and PCZ had one of the highest synergy scores and efficacy effects in LU-NB-1 and LU-NB-2 models both at day 3 and day 7 (**Fig. 2B,C and Fig. S2B**). In addition, PIT had the highest bioavailability and favorable pharmacokinetics/pharmacodynamics of the statins(*31*).

The cell death effects of the treatment were confirmed visually and by PI staining (**Fig. 2D,E**). Overall, treatment of chemoresistant NB organoids with statins in combination with phenothiazines resulted in strong synergy and cell death.

### Combined prochlorperazine and pitavastatin is effective in an NB PDX model

We next tested PCZ and PIT alone and in combination *in vivo* (**Fig. S2C**). Mice were randomly allocated to treatment groups when tumors reached at least 150 mm^3^. Mice were treated intraperitoneally (i.p.) with PCZ, PIT, and their combination (5 mg/kg each or drug vehicle daily, six times a week for six weeks) (**Fig. S2C**). The average tumor size at the start of treatment was ∼200 mm^3^ (**Fig. S2D**). Treatment was well tolerated, with only transient drowsiness seen with PCZ and no evidence of weight loss (**Fig. S2E**). However, i.p. administration of the drugs did not reduce tumor growth (**Fig. S2F**), and transcriptomic analysis of excised tumors did not reveal any group-specific clustering (**Fig. S2G**). Given these results, it was likely that i.p. delivery could not achieve effective drug concentrations in the tumors, possibly due to first-pass metabolism(*32*).

We therefore moved forward using direct administration of the drugs (alone or in combination) into the tumor by intratumoral (i.t.) injection in the treatment-resistant and aggressive NB model LU-NB-1 (**Fig. 2F**). The LU-NB-1 model is resistant to standard NB chemotherapy treatment *in vitro* and *in vivo*(*19*), reflecting the clinical scenario of a lack of response. LU-NB-1 PDX cells were injected subcutaneously into NSG mice, which were randomly allocated to control, single drug treatment, or combination treatment groups when the tumors reached over 350 mm^3^ (**Fig. S2H**). Mice were treated i.t. daily five times a week according to the group assignment (**Fig. 2F**) and were monitored for weight loss (**Fig. S2I**), tumor growth and survival (**Fig. 2G, Fig. S2J, K)**. Compared with control, treatment with the drug combination significantly delayed tumor growth (Dunnet’s multiple comparison test, p=0.023) at day 8 of treatment and prolonged survival of these mice (p=0.001, log-rank test between controls and combination) (**Fig. 2G, H**). Thus, while systemic administration of the PIT-PCZ drug combination did not affect tumor growth, intratumoral treatment decreased tumor growth and prolonged survival in a chemoresistant NB PDX model.

### Combined prochlorperazine and pitavastatin modulates cholesterol synthesis pathways

We performed transcriptomic analysis by RNA-seq to investigate the molecular mechanisms of action of the drug combination *in vitro*. After 48 h, PIT and the combination treatment induced the greatest differential gene expression compared with controls, with PCZ producing more subtle changes (**Fig. S3A**). As expected from the previously described drug mechanisms, GSEA suggested that PCZ inhibits dopaminergic synapse responses, whereas PIT modulates cholesterol metabolism (**Fig. 3a, Fig. S3B**). Unexpectedly, PCZ treatment also increased expression of genes involved in cholesterol and steroid metabolism. Both drugs were associated with downregulation of genes involved in DNA replication (**Fig. 3A, Fig. S3B**). Directed analysis of KEGG metabolic pathways indicated that several biosynthesis processes, including the steroid and terpenoid backbone and lipid metabolism, were strongly upregulated (**Fig. 3B**). Downregulation of drug and nucleotide metabolism and oxidative phosphorylation were consistent with a reduction in normal proliferation (**Fig. 3B**).

**Fig. 3.**
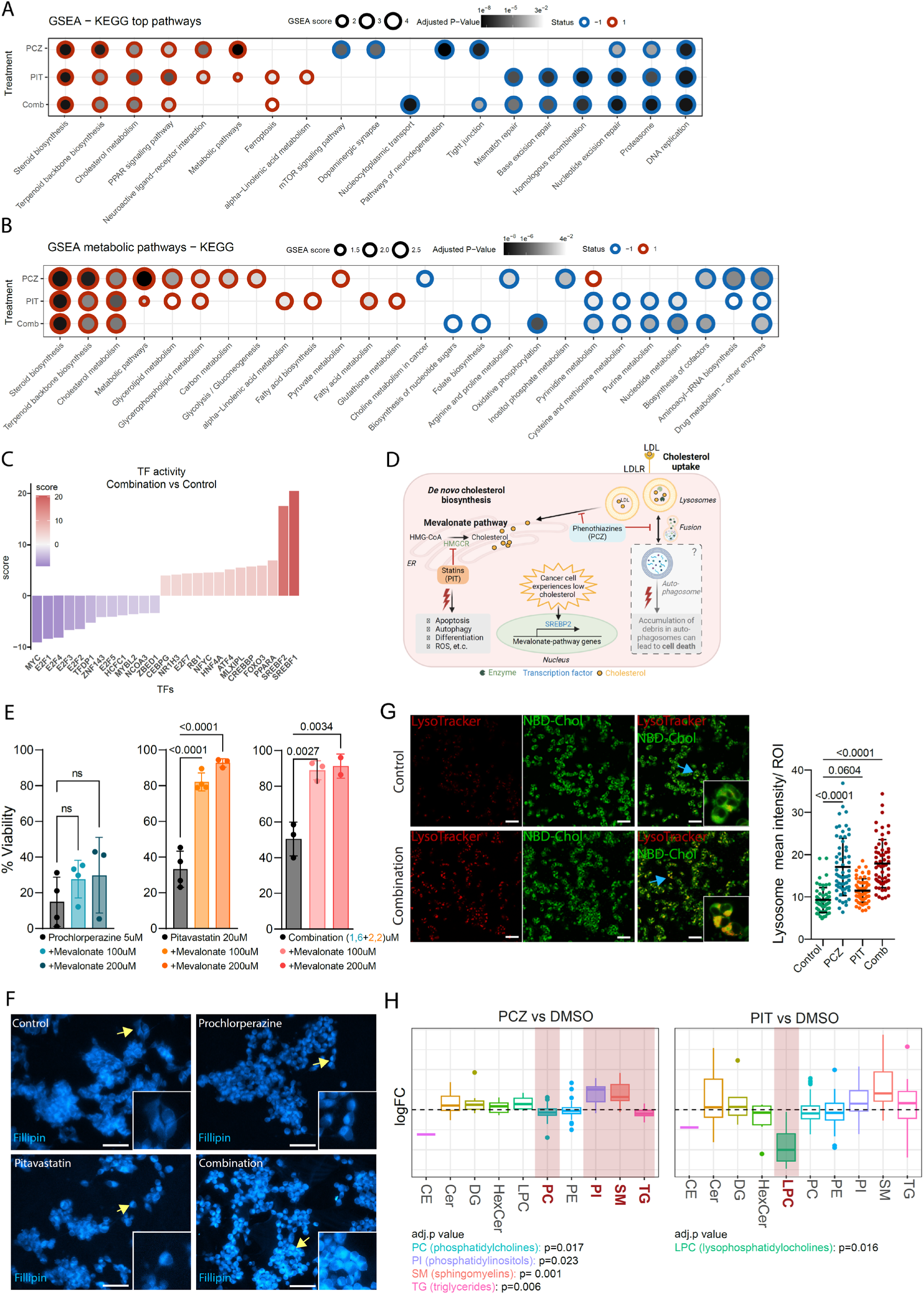
Dysregulated cholesterol metabolism after PCZ+PIT treatment. **A** GSEA transcriptomic analysis of treatment response in LU-NB-2 PDX cells after 48 h treatment using KEGG pathways. **B** Directed GSEA analysis of KEGG metabolic pathways. **C** Transcription factor (TF) activity inferred with CollecTRI after combination treatment. **D** Schematic overview of a potential dual-hit mechanism of action for PCZ and PIT in cholesterol uptake and biosynthesis. **E** Mevalonate rescue experiments after treatment with PIT, PCZ, and combination in LU-NB-2 organoids (one-way ANOVA followed by Dunnet’s multiple comparisons). **F** Representative photos of treated LU-NB-2 cells stained with the free-cholesterol binding dye filipin (scale bar 50 µm). **G** Live cell imaging using LysoTracker staining and labelled cholesterol (scale bar 50 µm) followed by quantification of mean lysosome intensity in 20 selected ROIs (cells); 24 h (one-way ANOVA with Tukey’s multiple comparisons test). **H** Lipidomics analysis showing changes in abundance of lipid classes in PCZ and PIT treatment groups compared with DMSO-treated controls; significant changes marked in red (permutation test was used to calculate p-values and corrected with Benjamini-Hochberg test, adj.p-values: PC:0.017, PI:0.023, SM:0.001, TG:0.006, LPC:0.016). GSEA-gene set enrichment analysis; PCZ-prochlorperazine; PIT-pitavastatin; TF-transcription factor; LDL-low density lipoprotein; LDLR-low density lipoprotein receptor; CE-cholesterol esters; Cer-ceramides; DG-diacylglycerol; HexCer-hexosylceramide; LPC-lysophosphatidylcholine; PC-phosphatidylcholine; PE-phosphatidylethanolamine; PI-phosphatidylinositol; SM-sphingomyelin; TG-triglycerides.

Inference of transcription factor (TF) activity after drug combination treatment showed low MYC and E2F activity, consistent with decreased NB cell proliferation (**Fig. 3C**). This was further supported by observed downregulation of phosphorylated Akt protein after combination treatment (**Fig. S3C**). Both single and combined drug treatment resulted in striking upregulation of SREBF1 and SREBF2, TFs essential for cholesterol synthesis (**Fig. 3C, Fig. S3D**). This upregulation could be explained by a compensatory effect in PIT-treated cells with statin-induced inhibition of the rate-limiting HMGCR enzyme in the mevalonate pathway, downstream of SREBF (**Fig. 3D**). Rescue experiments to reverse the effect of PIT with mevalonate, a product of the HMGCR reaction, confirmed this causation (**Fig. 3E**). However, mevalonate did not alter the viability of PCZ-treated cells, suggesting that a different mechanism was responsible for reduced viability and upregulation of factors influencing cholesterol synthesis (**Fig. 3D, E**).

Phenothiazines (e.g., PCZ) are reported to exert multiple effects that might alter proliferation and cell death, including inducing apoptosis, G1 cell cycle arrest, and membrane destabilization. Our transcriptomic data suggested that PCZ led to dysfunctional cholesterol metabolism (**Fig. 3A,B, S3E**). To investigate cellular cholesterol abundance and localization during treatment, we stained free cholesterol with filipin and observed an increased and more localized signal after combination treatment (**Fig. 3F**). Live cell experiments with LysoTracker and labeled cholesterol revealed co-localization and accumulation of cholesterol in lysosomes after PCZ and combination treatment but not as pronounced with PIT treatment alone (**Fig. 3G**). Together, our results suggest that PCZ affects lysosome membrane fusion, leading to cholesterol accumulation in lysosomes and an overall cellular deficit in cholesterol (**Fig. 3D**). Thus, PIT-induced inhibition of *de novo* cholesterol synthesis combined with PCZ-induced cholesterol accumulation in lysosomes create a dual-hit mechanism targeting cholesterol availability in NB cells.

Treatment-induced metabolic profiles were further investigated using lipidomic analysis, which revealed significant changes in individual lipid species (**Fig. S4**). Specifically, there was increase in sphingomyelin lipid class after PCZ treatment which has been previously linked to phenothiazine’s cytotoxic effects(*33*) (**Fig. 3H)**. Decrease in lysophosphatidylcholines after PIT treatment could point to smaller fatty acids pool needed for cell proliferation (**Fig. 3H**)(*34, 35*). These changes consistent with the known effects of the drugs influence on lipid and cholesterol availability and function in the cancer cells(*35*).

In conclusion, treatment of NB organoids with PIT and/or PCZ decreased proliferation and caused compensatory upregulation of cholesterol production through high transcriptional activity of transcription factors SREBF1/2, cholesterol cellular deficit, and cell death.

### Combined prochlorperazine and pitavastatin leads to a chemosensitive NB cell state in vitro

We next investigated the effect of the drug combination on NB cell states *in vitro*. Given that different gene sets can define differentiated ADR and undifferentiated MES–like cell states, we explored multiple signatures derived from *in vitro* models, xenografts, and patient tumors (van Groningen(*12*), Boeva(*13*), Gartlgruber(*16*), Olsen(*18*), Manas(*19*), Bedoya-Reina(*15*), Yuan(*17*), Thirant(*21*), and Patel(*22*); **Table S4**). GSEA showed enrichment of ADR gene signatures derived from patient tumors (Bedoya-Reina, Yuan, and Patel) and an integrated signature from multiple datasets (Manas). This analysis indicates that PCZ-PIT combination treatment increased ADR-like signatures (**Fig. 4A, B**) and decreased undifferentiated MES-like signatures (**Fig. 4A, C**). The strong decrease in SYMP-Patel signatures likely reflects the decreased NB cell proliferation following treatment.

**Fig. 4.**
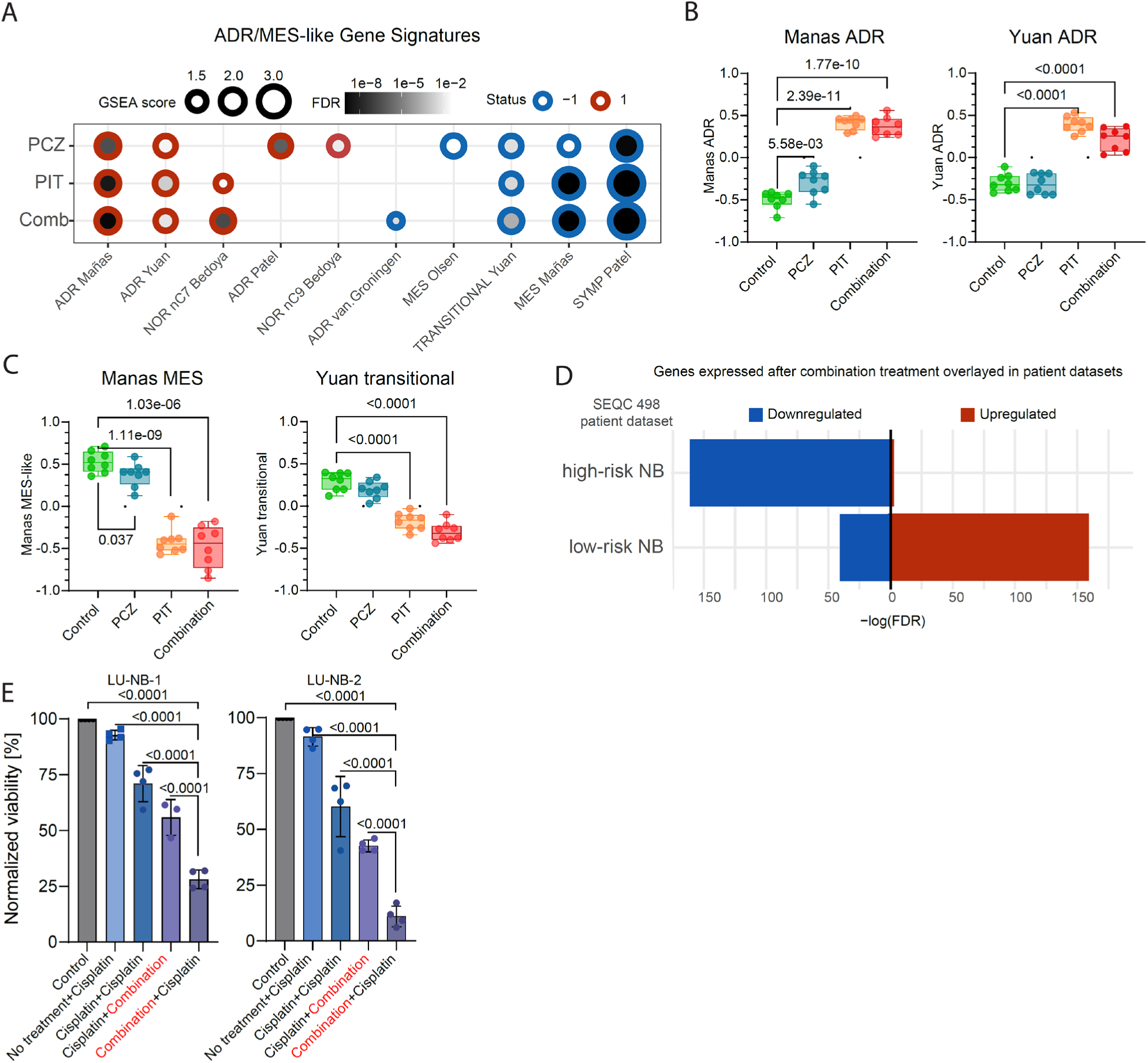
Phenotypic shift after PCZ+PIT treatment *in vitro.* **A** Association between various MES/ADR-like gene signatures and DEGs in LU-NB-2 organoids treated with PCZ and/or PIT for 48 h. Only significant (FDR p-value<0.05) results are displayed. **B** Comparison between treatment groups for two individual ADR-like gene sets (Welch t-test). **C** Comparison between treatment groups for the Manas MES-like and Yuan transitional gene signatures (Welch t-test). **D** Association of treatment-induced gene expression with high-risk and low-risk NB features obtained from public patient dataset SEQC498. **E** NB cell viability after various sequential treatment strategies of the PCZ+PIT drug combination and cisplatin (one-way ANOVA with Tukey’s multiple comparisons test). PCZ-prochlorperazine; PIT-pitavastatin; Combination-PCZ+PIT; MES-like-undifferentiated mesenchymal-like cells; ADR–adrenergic/noradrenergic-like cells.

We further investigated the relationship between PCZ-PIT combination-induced phenotypes and those found in NB patient tumors. Using publicly available NB datasets (SEQC 498 and 161 Target), we found that genes upregulated after PCZ-PIT treatment were upregulated in low-risk NB tumors, whereas treatment-induced downregulated genes were representative of high-risk NB tumors (**Fig. 4D, Fig. S5A, B**). Additionally, we analyzed the datasets originally included in the *in silico* predictions (**Table S1**) and concluded that the transcriptional responses to PCZ and/or PIT were indeed associated with lower risk/stage groups (**Fig. S5C**).

Based on the drug-induced increase in an ADR signature, we hypothesized that pretreatment with the drug combination would sensitize NB cells to chemotherapy. Thus, we tested the combined PCZ-PIT treatment together with cisplatin in treatment-resistant NB organoids, LU-NB-1 and LU-NB-2. As predicted, different treatment sequences revealed that pretreatment with combined PCZ-PIT followed by cisplatin resulted in the lowest NB cell viability (**Fig. 4E, Fig. S5D**), suggesting that PCZ-PIT combination pretreatment can sensitize chemoresistant NB to chemotherapy.

Together, our data suggests that treatment with PCZ-PIT leads to a phenotype shift towards the ADR cell state *in vitro* and features of low-risk NB. Sequential treatment with PCZ-PIT followed by chemotherapy, sensitizes resistant NB cells in vitro to chemotherapy.

### Combined prochlorperazine and pitavastatin sensitizes chemoresistant NB to chemotherapy in vivo

Encouraged by the *in vitro* findings, we evaluated the effect of combined PCZ-PIT with standard-of-care COJEC chemotherapy *in vivo*. We used the resistant LU-NB-1 PDX model, which displays a very limited response to COJEC induction therapy(*19*).

In a short-term *in vivo* study for 10 days (**Fig. 5A**), both the drug combination and the combination + COJEC inhibited NB tumor growth (p=0.0045 and p<0.0001, respectively; mixed effects analysis with Dunnett’s multiple comparison test; **Fig. 5A,B**), with the triple combination being clearly more effective. The tumors had a mean starting volume of ∼300 mm^3^, and no toxicity and weight loss occurred over the duration of the study (**Fig. S6A**-**B**). To investigate the mechanisms of treatment response at the transcriptional level, we performed RNA-seq of tumors after 10 days of treatment. This analysis revealed upregulation of diverse stress responses and cholesterol homeostasis and downregulation of oxidative phosphorylation and MYC targets in the treated groups (**Fig. 5C)**. Transcriptional phenotype changes included upregulation of both the ADRN and MES-like cell states, possibly reflecting contribution of the complex tumor microenvironment *in vivo* and/or development of resistance (**Fig. 5D**).

**Fig. 5.**
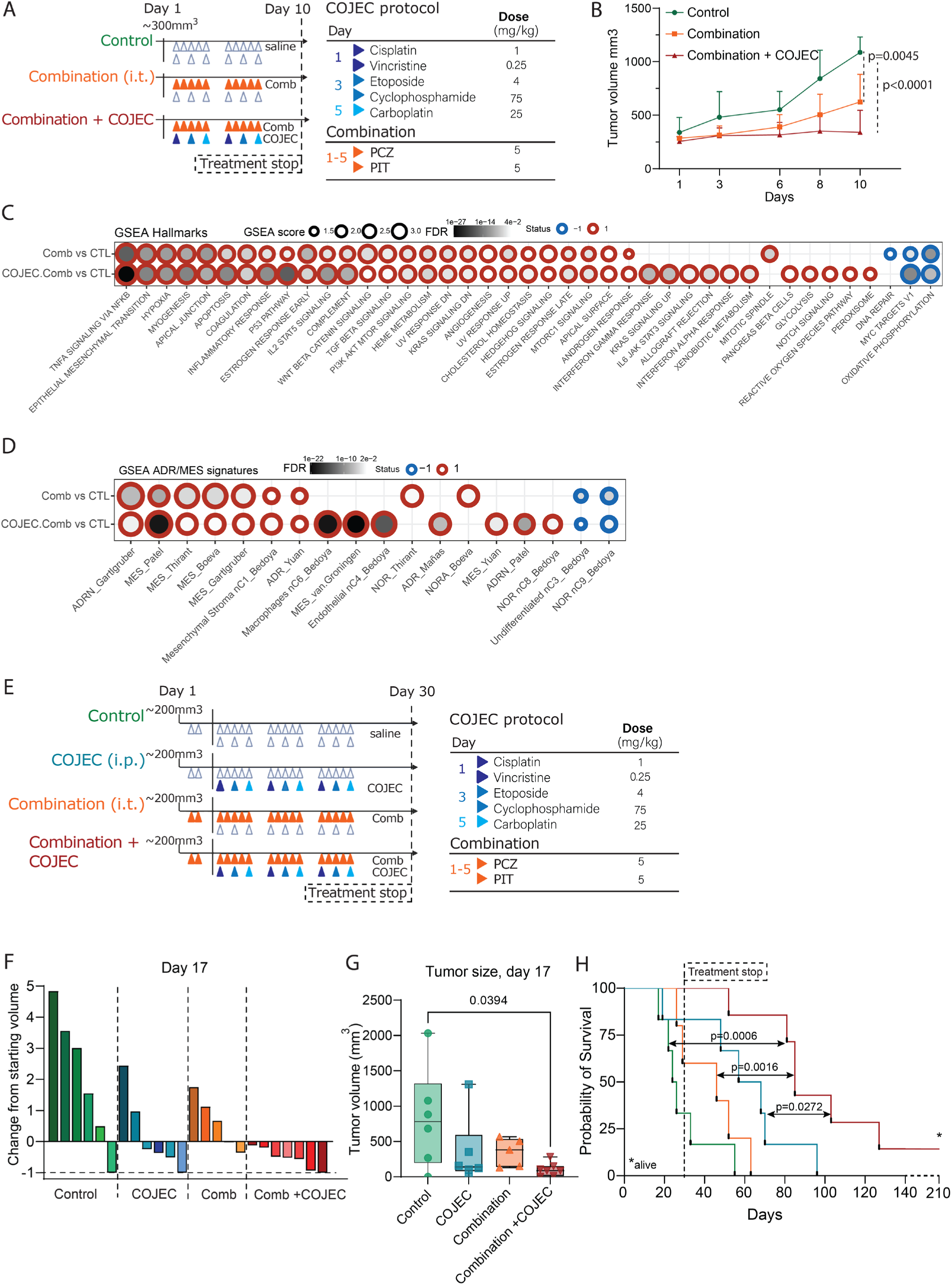
Addition of PCZ+PIT combination to standard-of-care COJEC outperforms COJEC treatment alone. **A** Schematic overview of short-term *in vivo* study of the combination PCZ+PIT (i.t.) with COJEC (i.p.). Mice with LU-NB-1 tumors were randomly allocated to the groups and treated with the corresponding schedules for 10 days. **B** Change in tumor volume over time for all groups (mixed effects analysis with Dunnett’s multiple comparison test). **C** RNA-seq and gene set enrichment analysis of 20 tumors (control=7, combination=7, combination + COJEC=6). Significant contrasts in expression between the two treatment groups and control are displayed for Hallmarks **D** Enrichment of MES- and ADR-like gene signatures after *in vivo* treatment. **E** Schematic overview of the long-term survival study in mice bearing LU-NB-1 tumors. Two doses of the combination were given before the start of standard-of-care COJEC chemotherapy. Treatment was given for 30 days. **F** Day 17 (last day of all mice alive) single mouse results showing % of tumor size change from baseline. **G** Summary of day 17 tumor volume (Tukey’s multiple comparison test of all groups). **H** Kaplan-Meier survival graph showing survival of mice treated with combination + COJEC compared with control (log-rank test p=0.006) or standard-of-care (log-rank test p=0.0272). PCZ-prochlorperazine; PIT-pitavastatin; combination-PCZ+PIT; i.t.-intra-tumoral; GSEA-gene set enrichment analysis; MES-like-undifferentiated mesenchymal-like cells; ADR– adrenergic/noradrenergic-like cells.

Finally, we compared PCZ-PIT combination + COJEC and COJEC treatment alone in a long-term survival study of treatment-resistant LU-NB-1 xenografts. Mice were randomized into four groups, and the treatment was initiated when tumors reached 200-300 mm^3^ (**Fig. 5E, Fig. S6C**). All tumors that reached that size were pre-treated with two i.t. injectionsof the combination before the i.p COJEC treatment started (**Fig. 5E)**. Intra-tumoral drug injections were performed according to a standardized schedule to ensure good drug distribution into the tumor (**Fig. S6D**). The addition of PCZ-PIT did not increase the side effects expected from COJEC chemotherapy (**Fig. S6E**). Mice were monitored for toxicity and showed <10% weight loss from baseline in the treatment groups that included chemotherapy (**Fig. S6E**).

Treatment with PCZ-PIT combination + COJEC decreased tumor growth compared with controls at day 17 (p=0.0394; Tukey’s multiple comparison test of all groups; the last day of all mice alive) (**Fig. 5F,G, Fig. S6F**). Tumor regression was observed in one control mouse, likely caused by the i.t. injections. A fraction of mice responded to combination or COJEC treatment alone, but tumors rapidly progressed after treatment was stopped at day 30 (**Fig. S6F**). Tumors in three of the six COJEC-treated mice initially regressed to 0 mm^3^ but quickly relapsed after treatment removal, with a median group survival of 62.5 days (range 19-96 days; **Fig. 5H, Fig. S6F**). In the combination + COJEC group, six out of seven tumors initially regressed completely (**Fig. 5F, Fig. S6F**), and survival was extended to a median of 85 days (range 52 to 210 days; **Fig. 5H**). One mouse in the combination + COJEC group did not relapse throughout the study period (210 days). Importantly, survival was significantly prolonged in the combination + COJEC group compared with both the control (p=0.006) and COJEC-treated mice (p=0.027, two-sided log-rank tests; **Fig. 5H**). This result underscores the benefit of adding the combination to standard-of-care chemotherapy.

In summary, the addition of PCZ-PIT combined therapy to standard-of-care COJEC chemotherapy outperforms COJEC chemotherapy alone and significantly prolongs survival in a chemoresistant NB PDX model.

## DISCUSSION

Chemoresistant high-risk NB is a major challenge in pediatric oncology, and novel treatment strategies are a clinical imperative. Through a drug repurposing approach, here we identified a novel combination of statins and phenothiazines (dopamine antagonists) with strong anti-NB properties. Exploiting DGEM based on multiple gene expression profiles in combination with the PRISM drug prediction tool, we identified effective and safe new therapies for high-risk NB. The novel drug combination was synergistic, altering cholesterol metabolism through a dual-hit mechanism and causing phenotypic switching of NB cells towards an ADR cell state, sensitizing them to chemotherapy. Treatment with a combination of PCZ and PIT together with standard-of-care COJEC chemotherapy outperformed chemotherapy alone and prolonged survival in a chemoresistant *MYCN*-amplified PDX model.

Drug repurposing has the potential to accelerate the long and expensive path from drug discovery to a clinically useful medication. By using already available safety, efficacy, and pharmacokinetic data, this approach is of particular interest for childhood and rare diseases where clinical trials are difficult to perform(*25, 36*). Diverse strategies have been applied to identify drugs for repurposing, including the Drug Repurposing Hub at Broad Institute(*37*), TargetTranslator(*38*), and DGEM(*23*). In this study we used two tools to predict potential treatments; the PRISM “guilt by association” tool, and the DGEM gene expression matching tool. Between these, many thousands of comparisons between diseases and compounds were carried out to arrive at a set of compound treatment predictions.

Twelve drugs were identified *in silico* for repurposing in high-risk NB, and subsequent experiments narrowed the list down to two classes of therapeutics: cholesterol synthesis inhibitors (statins) and anti-psychotic dopamine receptor (D2) antagonists (phenothiazines). Statins inhibit HMG-CoA reductase which reduces cholesterol production via inhibition of the mevalonate pathway (**Fig. 3D**) Statins are best known for reducing the risk of cardiovascular disease, however the impact of these drugs in cancer has been investigated for decades, and experimental studies in NB suggest that statins can induce apoptosis and differentiation(*38–40*). The antipsychotic phenothiazines, e.g. PCZ, are also indicated for use against nausea and migraine during chemotherapy treatment(*41*). Here we identified several phenothiazines (TFP, TRD, and PCZ) with potential anti-NB effects, and we selected PCZ as the most promising agent. Importantly, combined PCZ+PIT therapy yielded substantial synergistic effects on high-risk NB organoids.

As expected, gene expression and mechanistic analyses suggested that the mechanism of action of PIT was related to the mevalonate pathway(*31*). Consequently, the inhibitory effect on viability was rescued by mevalonate. Conversely, even if we cannot exclude that downregulation of dopaminergic activity is of partial importance for the antitumor effect of PCZ, this mechanism does not seem to be the main mode of action. Instead, the dominant pattern of treatment response after PCZ was strong upregulation of cholesterol biosynthesis and metabolism, similar to recent findings described in glioblastoma after treatment with phenothiazines(*42*). Cholesterol metabolism is known to be dysregulated in cancer, since both synthesis and uptake are often increased to meet the metabolic needs of cancer cells(*43*) (**Fig. 3D**). While statins are known to inhibit *de novo* cholesterol production, our data from live cell imaging showed accumulation of cholesterol and co-localization of cholesterol and lysosomes after treatment with combined PCZ+PIT. Thus, our data suggests that PCZ+PIT treatment targets cholesterol in cancer through a dual-hit mechanism of cholesterol accumulation in lysosomes (PCZ) and inhibition of *de novo* cholesterol synthesis (PIT). This finding is consistent with recent studies suggesting that phenothiazines can inhibit lysosome function, leading to impaired intracellular delivery of LDL-derived cholesterol(*44*). It has been suggested that phenothiazine-induced impairment of lysosome function can block lysosomes and autophagosome fusion, also leading to autophagy inhibition(*45*). In turn, dysfunctional autophagy can lead to the accumulation of toxic products such as ROS and oncogenic proteins in autophagosomes, causing the eventual death of NB cells(*46*).

The PCZ+PIT combination also led to a phenotypic shift *in vitro*, where tumor cells became more ADR-like and less MES-like. It has previously been suggested that MES-like NB cells contribute to NB chemotherapy resistance(*12, 19*), so we hypothesized that the drug-induced phenotypic switch would sensitize NB to chemotherapy. Indeed, using a chemoresistant PDX-model and a clinically relevant chemotherapy treatment protocol(*19*), we showed that mice treated with both the drug combination and COJEC standard-of-care chemotherapy displayed more frequent tumor regression and longer survival compared with standard-of-care treatment alone. Recent findings in small cell lung cancer (SCLC) have shown that statin treatment can sensitize tumors to chemotherapy to produce durable responses in a pilot study of patients with relapsed SCLC after chemotherapy(*47*), supporting this approach to treat chemoresistant and relapsed cancer. Suggested molecular mechanisms explaining increased chemosensitivity upon cholesterol deregulation include destroyed lipid rafts leading to aborted multi-drug resistant (MDR) ABC transporter activity(*48–51*) as well as downregulation of pathways of importance for epithelial-to-mesenchymal (EMT) cell state transition (e.g., Wnt, TGF-β)(*43*).

This study has some limitations. The generalizability of our study is limited as only *MYCN*-amplified PDX models were tested, and the effects of the combination in non-*MYCN* amplified high-risk NB is unknown. However, patient datasets used for the drug predictions were derived from both *MYCN* and non-*MYCN* amplified NB, suggesting that the drug combination could be effective in non-*MYCN* amplified NB as well. One challenging aspect of the novel combination of statins and phenothiazines is the bioavailability of the drugs in oral form(*31, 32, 52*). Intraperitoneal injection of the drugs did not produce significant effects as statins are known to undergo a first-pass metabolism that results in liver accumulation of the drug. Pitavastatin, despite having highest bioavailability and different metabolic fate than other statins, is still preferentially distributed to liver and intestines, and it is likely the reason for the drug to not circulate at biologically relevant concentrations(*53*). Despite evidence that mevalonate and cholesterol are important in cancer(*48*), no selective targeting drugs are yet available. In the present study, anti-NB effects were obtained by intratumoral injection, which would be challenging in a clinical setting in patients with multiple distant metastases, so a new formulation is necessary to improve tumor exposure. Importantly, the safety of novel combinations within a new target group, e.g., children, must also be considered afresh(*36, 54*). Nevertheless, there are successful examples of repurposing adult drugs for pediatric patients in other diseases; for example, fenfluramine, developed in the 60s as an appetite suppressant, has been shown to be effective in Dravet syndrome, a rare neurological disease of infancy, and the FDA approved fenfluramine for this use in 2020(*55, 56*).

Pitavastatin and other statins are currently being investigated in clinical trials to assess their bioavailability in glioblastoma (NCT05977738) and their effects as part of combination treatment in patients with leukemia (NCT04512105) and breast cancer (NCT03454529). Prochlorperazine is FDA-approved for indications including nausea and vomiting in a post-chemotherapy, post-operative setting and off-label use to treat acute migraines in adults and children. These antiemetic effects could be advantageous in the treatment of children with cancer taking chemotherapy(*57–59*). Recent findings suggest that PCZ can also enhance the anti-tumor effects of monoclonal antibody therapies, possibly through inhibition of endocytosis(*60*). Consequently, there are ongoing clinical trials of high doses of PCZ in combination with paclitaxel, trastuzumab, and pertuzumab for HER2-positive metastatic cancer(*61*).

In summary, using *in silico* drug predictions and experimental drug testing in patient-derived tumor models, we show that transcriptomics and machine learning based drugs-disease matching is a viable option to identify new treatment strategies against NB. We identified two new classes of drugs which, when administered together, hold promise for patients with chemoresistant NB (**Fig. 6**). With increasing numbers of single-cell transcriptional profiles of aggressive resistant tumors and rapidly developing deep learning methods, we expect that this drug repurposing approach will yield additional promising therapeutic strategies against treatment-resistant cancers.

**Fig. 6.**
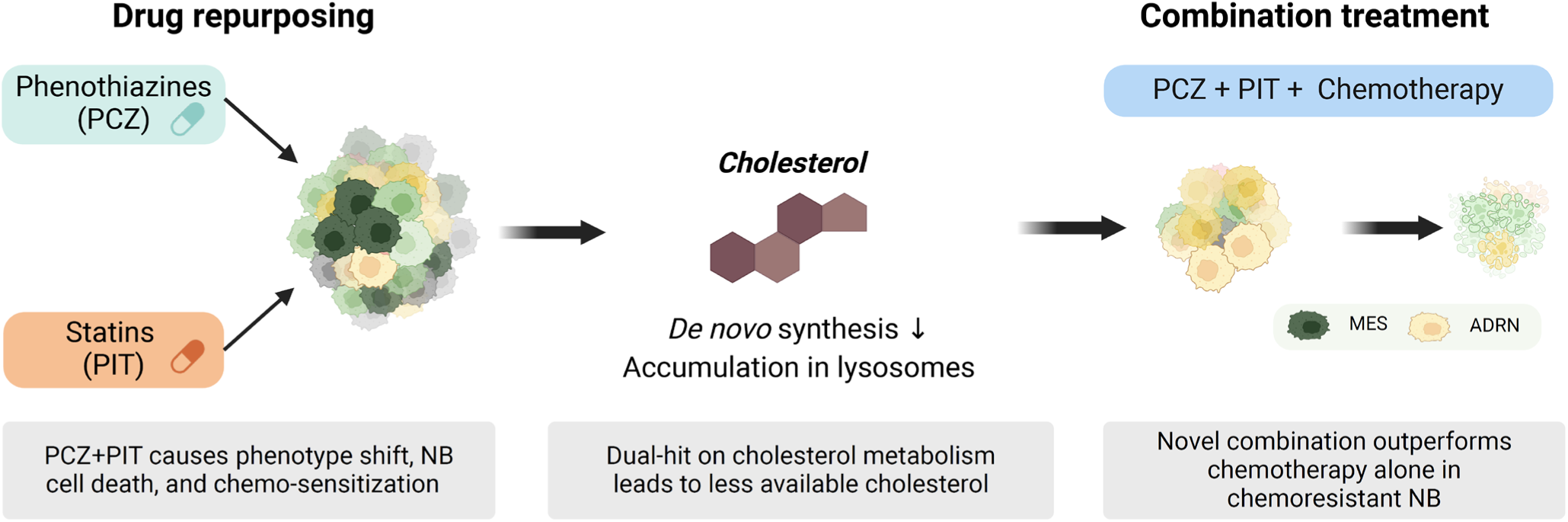
Schematic overview of suggested treatment effects after repurposed drugs combination together with standard-of-care chemotherapy.

## MATERIALS AND METHODS

### Experimental design

The objective of this study was to identify and test approved drugs for repurposing in high-risk neuroblastoma. We used a combination of *in silico* prediction methods (disease-gene expression matching and machine learning), *in vitro* experiments with neuroblastoma PDX-derived organoids, and *in* vivo high-risk neuroblastoma PDX models to investigate the effects of these drugs. For *in vitro* experiments, two or three biological replicates were performed (reported for each experiment). For *in vivo* studies, dissociated PDX organoids were injected into NOD scid gamma (NSG) mice (RRID:IMSR_JAX:005557) and upon tumor engraftment, mice were randomly assigned to control or treatment groups with a minimum of five mice per group. Mice were euthanized on the basis of tumor size, weight loss, overall health deterioration, or end of study time. Tumor tissues were collected from all mice for further analysis. All results were subjected to statistical analysis, with significance defined as p < 0.05. Investigators were not blinded to the experimental conditions, and no animals or data points in experiments were excluded from the analysis.

### In silico analyses for drug repurposing

Publicly available raw gene expression datasets were manually curated and analyzed for their suitability as input into Healx’s predictive technologies. Four raw gene expression datasets passed quality control criteria that ensured internal robustness of the input data (**Table S1**). Gene expression datasets were derived from samples obtained from patients at various stages of progression of tumor growth or who had died of the disease. NCBI GEO datasets were used to derive the NB disease gene expression signatures.

### Differential gene expression profiles (GEPs) and pathway enrichment analysis

NB GEPs were calculated based on differentially expressed genes between groups of progressive/severe outcome and successful outcome (details in **Table S1**) using an in-house bioinformatics workflow. Briefly, raw microarray datasets were downloaded from NCBI GEO, normalized by MAS5 or RMA, and subject to arrayQualityMetrics as defined by the curated contrasts to identify outliers. Datasets were supplied pre-processed. Outlier samples were removed from the contrasts, and differential gene expression analysis was performed using Limma (RRID:SCR_010943). Adjusted p-values were calculated according to the Benjamini-Hochberg procedure to test for false discovery, after removing insignificant (adj.P.Val > 0.05) and control probe sets from the signature. Gene expression profiles (GEPs) formed from differentially expressed genes in these datasets were evaluated with gene set enrichment analysis (GSEA) to ensure they described biological processes indicative of an NB phenotype (**Fig. S1A**).

### Drug matching

#### I. Disease-Gene Expression Matching (DGEM)

All GEPs passed our technical and biological quality assessments and were used to predict drugs with DGEM. Healx’s DGEM implements analysis algorithms that automatically select the set of conditions most likely to yield informative drug-disease connections. Briefly, known NB therapeutic agents serve as positive controls and are used to assess the performance of the DGEM experiment. Prior approved NB therapies or agents in late-stage clinical trials are assumed to be highly ranked in the experiment. The degree to which these positive control compounds are enriched provides a mechanism to optimize DGEM parameters and rank the prediction sets generated.

#### II. PRISM analysis

PRISM (RRID:SCR_005375) is a predictive methodology that utilizes known information about compound and disease similarities to infer novel associations between nearest-neighbor compounds and diseases. In detail, PRISM calculates high-dimensional feature vectors for each compound-disease pair. Similarly, to the repurposing method PreDR(*27*), PRISM employs a support vector machine (SVM) to identify clusters of known indication pairs within this high-dimensional space. We enhance the recall of this method by using a Kronecker product kernel in the SVM (*62*). Pairs of compounds and diseases found within or near these learned clusters are predicted as potential novel treatment indications. For instance, suppose compound B is known to treat disease A (**Fig. 1**). Compound D has a similar structure to compound B so, by association, is a predicted treatment for Disease C. Similarly, Disease C (lower right; red) is associated with disease A by phenotype. Hence, compound B and compound D may be appropriate treatments for disease C.

#### III. Literature mining

Healx employs literature mining and natural language processing to find compound/disease relationships in the scientific literature. Text mining was employed to identify mentions of compounds and diseases in the titles and abstracts of millions of scientific articles published from 2000 to 2016 and extracted sentences where both a compound and a disease occurred. We used deep learning methods to classify those sentences and generate therapeutic scores, predicting the likelihood that the compound could treat the disease. We previously validated this method and showed strong recall of known therapeutic relationships(*23*).

The evidence from this approach was automatically attached to predictions generated by other repurposing methods. Literature mining results were used as direct evidence for an association between a drug and a disease but also to infer evidence from related diseases and drugs. Literature mining is an important addition to known drug indications, as it can capture off-label treatment and preclinical experimental results.

#### IV. Treatment profiles

Treatment profiles specify the properties and medicinal qualities of a prospective drug. Healx referenced the NB treatment profile during the derivation of the recommended predictions.

### Drug repurposing recommendations

Healx performed an expert pharmacological review of the top-ranking drug predictions across each of the GEPs that were used as input to DGEM. Prediction results were first analyzed by domain experts at Healx, who provided annotations with supporting pharmacodynamic data from an internal knowledgebase relevant to inferring therapeutic benefit in NB, such as literature support, clinical trials, and drug mechanism of action. This produced a ranking and filtering of strongly predicted drugs that were likely to be most efficacious, such that a case may be made for prospective validation of each drug by a team of drug repurposing experts.

### Drugs

Fourteen small molecule drugs were purchased from Selleckchem (Houston, TX): trifluoperazine (S3201), thioridazine (S5563), nortriptyline (S3698), prochlorperazine (S4631), resveratrol (S1396), sildenafil (S1431), dasatinib (S1021), pazopanib (S3012), lovastatin (S2061), folic acid (S4605), topiramate (S1438), tianeptine (S5087), fluvastatin (S1909), and pitavastatin (S1759). Drugs were stored according to the manufacturer’s instructions. Stock concentrations were made by dilution in DMSO. Drugs were further diluted in cell culture medium for testing *in vitro*.

### Tumor organoid cultures

NB tumor organoids were established from PDX *in vivo* models and cultured as free-floating spheres in serum-free medium with the addition of epidermal growth factor (EGF) and basic fibroblast growth factor (bFGF) as described(*28, 63*) (**Table S2**). Models were verified by SNP analysis and regularly tested for *Mycoplasma*.

### Single drug testing cell viability assays

LU-NB-1, LU-NB-2, and LU-NB-3 PDX-derived tumor organoids were dissociated into single cells and seeded into opaque 96-well plates (Corning Inc., Corning, NY), 5000 cells per well, and treated with a range of seven concentrations (0.014 to 10 µM) of each of the drugs (**Table 1**). Cells were incubated for 3 or 7 days. Cell viability was normalized to control wells based on a metabolic readout with CellTiter-Glo (G7571; Promega, Madison, WI) luminescence. Luminescence was measured with a Synergy2 Multi-Mode plate reader (BioTek, Winooski, VT). Biological triplicates were used.

### Combination drug testing

Tumor organoids were dissociated into single cells and seeded into 96-well plates (5000 cells per well in a volume of 80 μl). The cells were treated with 10 µl of each statin and 10 μl of each antipsychotic at varying concentrations. Tumor organoids were incubated for 3 or 7 days and analyzed for cell viability using the CellTiter-Glo assay according to the manufacturer’s instructions. Cell viability matrices were used to calculate synergy scores and most synergistic area scores (MSA) using FIMM SynergyFinder (RRID:SCR_019318, https://synergyfinder.fimm.fi)(*64*). SynergyFinder compares the observed drug combination responses to the expected responses calculated using a synergy modeling method (here Zero Interaction Potency (ZIP) method)(*65*). The predicted response is compared with experimental data, and the synergy score (δ) is calculated as the percent of response beyond expectation. Biological duplicates were performed.

### Cell death assays

NB LU-NB-1 and LU-NB-2 cells (1 × 10^6^ cells/T25 flask) were treated with a ∼IC_20_ concentration of prochlorperazine (PCZ) and pitavastatin (PIT) or a control amount of DMSO. Cells were allowed to grow for 7 days, and photos were taken at the end of the treatment time with an Infinity 1 camera (10x) and the Infinity Analyze software (Lumenera, Ottawa, Canada). Further, treated organoids were dissociated to single cells using Accutase (#A6964, Sigma-Aldrich, St Louis, MO) and stained with propidium iodide (PI; P3566, Invitrogen, Waltham, MA). Flow cytometry was performed with a FACS Melody flow cytometer (BD Biosciences, Franklin Lakes, NJ), and the data were analyzed using FlowJo 10.8.1 software (RRID:SCR_008520). Biological duplicates were performed.

### In vivo study (intraperitoneal, i.p.)

All *in vivo* studies were conducted according to Regional Ethics Committee for Animal Research in Lund guidelines (ethical permit no. 19012-19). LU-NB-2 PDX tumor cells (2 × 10^6^) were suspended 3:1 in medium/Matrigel (Cat No.354234, Corning Inc., Corning, NY) and injected subcutaneously into the flanks of female NSG mice. When tumors reached >200 mm^3^, each mouse was allocated randomly to a group: control, PCZ, PIT, or combination (PCZ+PIT). Drugs were formulated at 1 mg/ml in 2.5% DMSO, 2.5% Tween 80, 40% PEG, 55% PBS. Mice were treated intraperitoneally (i.p.) with daily injections six times a week according to the group assignment (**Fig. S2C**). The control group was administered vehicle only, PCZ and PIT were given at a dose of 5 mg/kg, and the combination group received two injections, one of each drug. Tumor volumes in mm^3^ were measured with calipers and calculated according to the formula *V* = π(length)(width)^2^. The weight of the mice was monitored three times a week throughout the study. The study was finished on day 33 of treatment or earlier if the tumor exceeded 1500 mm^3^. Mice were sacrificed, and tumor pieces were divided and snap-frozen or fixed in 4% paraformaldehyde and embedded in paraffin.

RNA was extracted from snap-frozen tumor pieces in the i.p. study utilizing the AllPrep DNA/RNA Mini Kit (Qiagen, Hilden, Germany). The mRNA library preparation phase was executed employing the TruSeq Stranded mRNA Library Prep kit (20020594, Illumina, San Diego, CA). Clean up steps were automized with King Fisher FLEX (18-5400620, Thermo Fisher Scientific, Waltham, MA), and incubations and PCR were with the Eppendorf Mastercycler X50s (6311000010, Eppendorf, Hamburg, Germany). Subsequently, sequencing was carried out using the NovaSeq 6000 System (20012850, Illumina) with the NovaSeq 6000 S1 Reagent Kit, 300 cycles v1.5 (20028317), Phix Control (FC-110-3001, Illumina). Reads were aligned using STAR software, and the reference genome sequence was the Human GRCh38 primary assembly from the Ensembl database with annotation (GTF) from GENCODE v33 (RRID:SCR_002344, RRID:SCR_014966). Raw counts were normalized with DESeq2. The datasets were then uploaded onto the R2: Genomic Analysis and Visualization Platform (http://r2.amc.nl) under the designation “PDX Neuroblastoma P4-combo-in-VIVO_LUNB2_2022_117 - Aaltonen - 22 - custom - gencode33”. These procedures were performed by the Center for Translational Genomics, Lund University, and Clinical Genomics Lund, SciLifeLab. Analysis on the R2 platform investigated the top 100 most differentially expressed genes (ranked by SD, log_2_ transformed).

### In vivo study (intratumoral, i.t.)

LU-NB-1 PDX tumor cells (1 × 10^6^) were suspended in medium/Matrigel (3:1) and injected subcutaneously into the flanks of female NSG mice. When tumors reached >200 mm^3^, each mouse was allocated randomly to a group: control, PCZ, PIT, or combination. Drugs were formulated at 3 mg/ml and 6 mg/ml in 2.5% DMSO, 2.5% Tween 80, 40% PEG, 55% PBS. Mice were treated intratumorally (i.t.) with daily injections five times a week according to the group assignment (**Fig. 2F**). The control group was administered vehicle only (50 µl), whereas PIT and PCZ were given at a volume of 50 µl (c=3 mg/ml of the respective drug). In the combination group, mice received one injection of 25 µl PCZ + 25 µl PIT (c=6 mg/ml) mixed directly before administration. The dose of PIT and PCZ was ∼5 mg/kg, and the total volume of the injection was 50 µl. Tumor volumes in mm^3^ were measured with calipers and were calculated according to the formula *V* = π(length)(width)^2^. The weight of the mice was monitored three times a week throughout the study. The study was stopped on day 30 of treatment or earlier if the tumor exceeded 1500 mm^3^. Mice were sacrificed and tumor pieces were divided and snap-frozen or fixed in 4% paraformaldehyde and embedded in paraffin.

### RNA sequencing in vitro

LU-NB-2 PDX cells were seeded in T25 flasks, allowed to grow for 24 h, and treated for 48 h with the combination of PCZ +PIT (1.6 µM and 6.7 µM, *n* = 6), PCZ only (1.6 µM, *n* = 6), PIT only (6.7 µM, *n* = 6), or DMSO (*n* = 6). The treatment resulted in a ∼15% decrease in NB cell viability in the combination sample (**Fig.S3F**). Cell pellets were collected, and RNA was extracted using the AllPrep DNA/RNA Mini Kit (Qiagen) and sequenced on a NovaSeq 6000 System (20,012,850, Illumina). Library preparation and mRNA sequencing were performed by the Center for Translational Genomic, Lund University.

### Quality control and bulk RNA-seq data pre-processing for in-vitro and in-vivo studies

Raw FASTQ files of the intro studies underwent quality control using the FastQC tool v0.12.1 (*66*) (RRID:SCR_014583) and were further filtered using the Trim Galore tool v0.6.10(*67*) (RRID:SCR_011847) with custom parameters: -q 25, –stringency of 3, and -l 50. After trimming, reads were pseudoaligned using Kallisto v0.48.0(*68*) (RRID:SCR_016582) in conjunction with the GRCh38 v109 *Homo sapiens* transcriptome from the Ensembl database(*69*). The transcriptome index was generated using both the cdna and ncRNA .fa files. MultiQC v1.14 (RRID:SCR_014982) was used to summarize the results of these steps(*70*). Pseudoalignment output files were further summarized to gene level using the *tximport* R package v1.30.0(*71*) with the corresponding annotation table (v109) from the *biomaRt* package v2.58.0(*72*) (RRID:SCR_019214). The gene-count matrix was imported into a DESeqDataSet object using DESeq2 v1.42.0(*73*) (RRID:SCR_000154) and assessed using the recommended DESeq2 vignette workflow, along with sample outlier detection proposed by Chen et al.(*74*). The in-vivo workflow followed a similar process, with the exception of the trimming tool. Fastp(*75*) was used for trimming with the following additional parameters: --trim_front1 3, --trim_front2 3, --trim_poly_g, --detect_adapter_for_pe, -- adapter_fasta, and --length_required 50. For index generation, the Ensembl GRCh38 v111(*76*) (Homo sapiens transcriptome release was used with the corresponding annotation table from biomaRt for summarization.

### Analysis of differential gene expression and gene set enrichment analysis (GSEA)

The DESeq function within DESeq2 was employed for differential gene expression (DGE) analysis, with no log_2_ fold-change threshold applied. The statistical analysis relied on the Wald test, and the reported p-values underwent adjustment using the Benjamini and Hochberg method. Log_2_ fold-change (log2FC) values were subjected to shrinkage using the apeglm method(*77*). Pairwise comparisons were conducted between samples treated with drug combinations and single drugs against non-treated samples. To streamline the full gene list for subsequent volcano plots and gene set enrichment analysis (GSEA), genes with missing values in the adjusted p-values were excluded. The resulting differentially-expressed genes were obtained for each contrast. Ranked files were generated by ordering the DGE table based on the log2FC column, serving as input for GSEA with the GSEA function from the ClusterProfiler v4.10.0 R package(*78*). Specific parameters were adjusted, including maxGSSize=2000, eps=0, and nPermSimple=10000. GSEA utilized MSigDB Hallmark gene sets from the msigdbr v7.5.1 R package(*79*). Additionally, the KEGG database (RRID:SCR_012773) was employed for GSEA using the gseKEGG function from ClusterProfiler. The rank list was converted to entrez symbols using the bitr function from the clusterProfiler package (RRID:SCR_016884). The enrichment results in **Fig. 3B** were manually curated to focus solely on pathways associated with metabolism. For the *in vivo* data, the differential gene expression analysis followed the same procedure. However, the ranked files for GSEA were generated using -log10(p-value) * sign(log2FC). Additionally, GSEA for the *in vivo* data was performed using only the MSigDB gene sets, without employing the gseKEGG function.

### Pathway and transcription factor activities inference

For transcription factor (TF) activity inference based on expression data, we leveraged the collecTRI gene regulatory network from the R package “decoupleR” v2.8.0, which encompasses a curated collection of TFs and their target genes(*80*). Activity was determined through a univariate linear model using the ulm function, with the top 25 TF values reported for each contrast. It is crucial to note a slight workflow variation in generating differentially-expressed genes for CollecTRI inference. Adjustments included introducing the parameter countsFromAbundance = “lengthScaledTPM” during gene-level summarization using tximport. Furthermore, genes with fewer than 4 log_2_ counts and those not expressed in at least 8 samples (minimum corresponding to a condition) were excluded. Genes lacking a gene symbol were also removed. The final step involved executing the differential gene expression analysis using the limma workflow(*81*), incorporating the lmFit, contrast.fit, and eBayes functions sequentially.

### Rescue experiments

LU-NB-2 PDX-derived tumor organoids were dissociated into single cells and seeded into opaque 96-well plates (Corning Inc.), 5000 cells per well, and treated with 20 µM of PIT, 5 µM of PCZ, and PIT + PCZ (6.7 + 5 µM, respectively) alone or with respective treatments with addition of mevalonate (100 µM and 200 µM). Cells were incubated for 3 days. Cell viability was normalized to control wells based on metabolic readout with CellTiter-Glo (G7571; Promega) luminescence. Luminescence was measured with a Synergy2 Multi-Mode plate reader (BioTek). Biological triplicates were used. Mevalonate solution was prepared fresh by dissolving mevalonolactone (<u>M4667</u>; Sigma-Aldrich) in 1 M HEPES buffer, and the pH was adjusted to ∼7 using 1 M NaOH solution. This was further diluted in medium, and pH was controlled to be neutral before addition to the cell culture.

### Western blotting

Cells were treated as in the *in vitro* RNA-seq experiment, and collected pellets were lysed in RIPA buffer and supplemented with complete protease inhibitor (Roche, Basel, Switzerland) and phosSTOP (Roche). Proteins were separated using SDS–polyacrylamide gel electrophoresis gels and transferred to nitrocellulose membranes (#1704271, Bio-Rad, Hercules, CA). Antibodies used were pAktSer(RRID:AB_331168, 193H12, Cell Signaling Technology, Danvers, MA), panAkt (RRID:AB_1147620, 40D4, Cell Signaling Technology), and actin (RRID: AB_2537667, **#**MA5-15739-HRP, Invitrogen) as a loading control. Phosphorylated and total proteins from the same biological replicate were probed on separate membranes with respective loading controls. Imaging was performed using Luminata Forte Western HRP substrate (MilliporeSigma, Burlington, MA) and Amersham Imager 600 (GE Healthcare Bio-Sciences AB, Chicago, IL). Three biological replicates were used.

### Filipin staining

LU-NB-2 cells were seeded on glass slides on laminin (10 μg/ml; LN521-05, BioLamina, Sundbyberg, Sweden). Cells were allowed to attach and were treated with PIT, PCZ, and PIT+PCZ or positive control U-18666A (ab133116, Abcam, Cambridge, UK). After 48 h treatment, cells were fixed in 4% paraformaldehyde and stained with filipin according to the protocol (ab133116, Abcam).

### Live cell imaging

To investigate cholesterol transport during drug inhibition, we performed live cell staining on LU-NB-2 cells using LysoTracker Red DND-99 (L7528, Invitrogen) and NBD-labeled free cholesterol (ab269448, Abcam). Cells, grown adhering to laminin (10 μg/ml; LN521-05, BioLamina), were loaded with LysoTracker, washed, labeled with cholesterol and treated accordingly. Cells were further imaged with a Zeiss confocal microscope (Carl Zeiss, Baden-Württemberg, Germany) from treatment to 24 h to observe the effects. Pictures for analysis were collected from three biological replicates at 24h time point. In each picture we identified 20 ROIs and measured red fluorescence intensity/ROI (Zen 3.1, Carl Zeiss, RRID:SCR_013672) and analyzed with GraphPad.

### Lipidomics analysis

LU-NB-2 PDX cells were seeded in T25 flasks (one million per flask), allowed to grow for 24 h, and treated for 48 h with the combination of PCZ + PIT (1.6 µM and 6.7 µM, *n* = 6), half of combination of PCZ + PIT, PCZ only (1.6 µM, *n* = 6), PIT only (6.7 µM, *n* = 6), DMSO (*n* = 6), cisplatin (5 µM), and the combination of PCZ + PIT with cisplatin. Cells were collected as pellets until extraction.

### Extraction

Lipids were extracted using the methyl tert-butyl ether (MTBE) extraction procedure(*82, 83*). A mixture of internal standards (IS) in methanol (150 µL) was added to each sample of one million cells. The following lipids were used as IS: LPC 12:0 (1 µg, Avanti Polar Lipids Inc., Alabaster, AL, US), PC 24:0 (0.5 µg, Larodan AB, Sweden), and TG 15:0-18:1-15:0-d7 (1 µg, Avanti Polar Lipids Inc., Alabaster, AL, US). Then, MTBE (750 µL) was added, and samples were vortexed for 5 min and sonicated for 5 min. Subsequently, 100 µL of water was added, and samples were vortexed for an additional 5 min, followed by sonication for 5 min and centrifugation for 10 min at 4°C at 10,000 rpm in an Eppendorf centrifuge 5424 R (Eppendorf). Next, 700 µL of the MTBE phase from each sample was transferred to new glass vials, and a quality control (QC) sample was prepared by mixing aliquots from all sample extracts. All samples and QC were then evaporated to dryness using an miVac Duo Concentrator (Genevac Ltd., Ipswich, UK) connected to a membrane vacuum pump (Welch by Gardner Denver, Ilmenau, Germany). Prior to LC/MS analysis, samples were reconstituted in an isopropanol/acetonitrile mixture (9:1, v/v, 50 µL) containing PC 15:0-18:1-d7 (0.5 µg per sample, Avanti Polar Lipids Inc.).

### Untargeted LC/MS lipid analysis

Lipidomics analysis was performed on an Agilent 1290 Infinity UHPLC system (Agilent Technologies, Santa Clara, CA) coupled with a timsTOF Pro 2 (Bruker Daltonics GmbH & Co. KG, Bremen, Germany). Chromatographic separation was performed on an Acquity UPLC CSH C18 column (1.7 µm, 2.1 x 75 mm; Waters Corporation, Milford, MA) equipped with an Acquity UPLC VanGuard CSH C18 pre-column (1.7 µm, 2.1 x 5 mm; Waters Corporation) as previously described(*83*). The mass spectrometer was operated in parallel accumulation serial fragmentation (PASEF) scan mode in the m/z range of 50-1500 m/z and 1/K_0_ range of 0.55-1.82 V·s/cm^2^. Electrospray ionization was performed in positive (ESI(+)) mode with end plate offset of 500 V, capillary voltage of 4500 V, nebulizer pressure of 2.2 bar, drying gas flow of 10 L/min, and drying temperature 220°C. During each sample acquisition, a calibration mixture was injected over 15-16 min. The calibration mixture contained Agilent Low Concentration Tuning Mix and a 10 mM sodium formate solution.

### Lipidomics data processing

Acquired data were processed in Bruker Compass MetaboScape 2022b software (v9.0.1, Bruker Daltonics GmbH & Co. KG, Bremen, Germany). Recursive feature extraction was applied to detect features if they were present in at least 3 out of 54 injected samples (including blanks and QC samples) and in at least 60% of one group corresponding to a cell treatment group. Peak intensity threshold was set to 1000 counts, with a minimum 4D peak size of 100 points. For ion deconvolution, the EIC correlation threshold was set to 0.8, and mass and mobility recalibration were performed for each sample. Peak annotation was performed based on the LipidBlast library(*84*) and the build-in rule-based lipid annotation tool in MetaboScape. Lipid species containing >20% missing data were excluded from subsequent analysis. Missing data were imputed using a conservative approach, setting them to half the lowest detected level. Following data preprocessing, the lipid dataset underwent quality control, multivariate analysis, normalization, differential analysis, and enrichment analysis. These analyses were carried out using the lipidR workflow(*85*). Notably, probabilistic quotient normalization (PQN) was employed for data normalization. Lipids showing significant differential abundance were identified using thresholds of p.adjusted value ≤5% and log_2_ fold-change ≥1. Enrichment analysis was performed by ranking the differentially abundant lipids based on their log_2_ fold-change values. For each predefined lipid set, an enrichment score (ES) was calculated using the fgsea package by assessing the overrepresentation of set members at the extremes of the ranked list. The significance of the enrichment scores was determined through permutation testing, generating a null distribution of ES values. P-values were adjusted for multiple testing to control the false discovery rate.

### Cell state enrichment analysis

To investigate transcriptional responses following treatment with drug combinations or single drugs, we curated a comprehensive gene set representing various cell states including differentiated adrenergic (ADR), immature mesenchymal-like (MES), sympathoblast (SYMP), Schwann cell precursor-like (SCP-like), and transitional states, drawing from established sources(*12, 13, 15–19, 21, 22*) (**Table S4**).

Additionally, we incorporated gene sets associated with different cell identities in low- and high-risk NB, as proposed by Bedoya-Reina et al.(*15*). Employing this expansive set of gene sets, we conducted GSEA using the rank files and the packages described in the “*Analysis of differential gene expression and gene set enrichment analysis (GSEA)*” section. It is noteworthy that, considering the limited number of genes comprising the SCP-like gene set, we adjusted the minGSSize parameter to 2 to ensure a meaningful analysis.

### In silico analyses of public patient cohorts

For bulk analysis, the SEQC cohort of 498 NB patients were used. Using this cohort, sets of genes significantly upregulated in patients classified into high- or non-high-risk groups and directly or inversely correlated with age at diagnosis were obtained. RPMs were log_2_-transformed, and the difference between risk groups was calculated for each gene using FDR-corrected ANOVAs. Pearson correlations were computed between gene expression and age at diagnosis with FDR-corrected p-values signaling the chances of obtaining the correlation coefficient in an uncorrelated dataset. Similarly, gene sets with significantly higher expression in poor or better survival cases were retrieved with the Kaplan–Meier Scaner Pro tool in R2, using FDR-correct p-values obtained with log-rank tests between groups of patients with different event free-survival probabilities. Significant genes were selected with an FDR threshold of 0.01. Gene enrichment was further calculated with one-sided Fisher’s exact tests corrected with the Benjamini–Hochberg approach. Using the same approach, genes significantly differentially expressed between ganglioneuroblastoma and NB were obtained for the TARGET cohort of 161 patients.

Raw microarray data files were obtained from GSE16237, GSE13136, GSE73537, and GSE3446. For the GSE3446 dataset, preprocessed normalized summarized experiments were utilized. Quality exploration of the raw unprocessed datasets was conducted using the arrayQualityMetrics package(*86*) (RRID:SCR_001335). Samples were selected based on specific criteria for each dataset (**Table S1**): GSE16237: *MYCN* amplified Stage 1, 2, 3, 4, and 4S; GSE13136: 11q deletions with non-*MYCN* amplification, 1p deletions with *MYCN* amplifications, and samples with no aberrations; GSE73537: *MYCN* amplified samples; GSE3446: primary tumor samples.

Normalization of each unprocessed raw dataset was performed using the RMA algorithm. Following normalization, intensity-based filtering was applied, removing transcripts with intensities below a manual threshold of 4 in at least as many arrays as the smallest experimental group. Additionally, probes with multiple mappings were excluded from the analysis. Differential expression analysis was conducted using the limma package (RRID:SCR_010943) (*81*), with design matrices defined as follows: GSE16237: *MYCN* amplified Stage 3, 4 (High Risk) vs. *MYCN* amplified Stage 1, 2, 4S (Low Risk); GSE13136: 11q deletions with non-*MYCN* amplification (High Risk 1) vs. samples with no aberrations (No Aberrations), and 1p deletions with *MYCN* amplifications (High Risk 2) vs. samples with no aberrations (No Aberrations); GSE73537: *MYCN* amplified samples Late Stage (LS) vs. *MYCN* amplified samples early stage (ES); GSE3446: primary tumor samples that relapse (TR) vs. primary tumor samples that do not relapse (TnR), with Seeger1 representing the U133A chip and Seeger2 representing the U133B chip. GSEA was performed using a ranked file generated by ordering the DGE table based on a ranking column, calculated as the product of -log10(p.value) * sign(logFC). The GSEA function from the ClusterProfiler v4.10.0 R package was employed, utilizing MSigDB Hallmark gene sets from the msigdbr v7.5.1 R package.

### In vitro treatment with PCZ-PIT combination together with chemotherapy

LU-NB-1 and LU-NB-2 PDX-derived tumor organoids were dissociated into single cells and seeded on laminin in opaque 96-well plates (Corning Inc.), 5000 cells per well, and treated with DMSO, PIT + PCZ, or cisplatin for 48 h. Then, media were exchanged to another treatment and incubated for another 48 h. Cell viability was measured with CellTiter-Glo (G7571; Promega) luminescence. Luminescence was measured with a Synergy2 Multi-Mode plate reader (BioTek). At least three biological replicates were used.

### In vivo study i.t.: combination + COJEC (short term)

LU-NB-1 PDX cells (1 x 10^6^) were injected subcutaneously into NSG females as above. When tumors reached 200-300 mm^3^, mice were randomized into treatment groups: control, PCZ + PIT combination, and PCZ + PIT combination + COJEC (**Fig. 5A**).

PCZ + PIT (total volume 50 µl; 5 mg/kg of each drug) was administered i.t. daily five times per week, as described for the i.t. study. The injection site was altered around the tumor to assure equal distribution across the tumor. COJEC treatment was given i.p. and consisted of five different chemotherapies distributed in cycles over one week (**Fig. 5A**); Day 1 – cisplatin + vincristine, Day 3 – etoposide + cyclophosphamide, Day 5 – carboplatin (see **Fig. 5A** for doses). All COJEC drugs were dissolved in saline. Treatment was administered for 10 days, after which tumor pieces were divided and snap-frozen or fixed in 4% paraformaldehyde and embedded in paraffin. Mouse weights and tumor volumes were measured three times per week throughout the study. RNA was extracted from snap-frozen tumor pieces in the short term i.p. study, and sequencing was conducted as above. These procedures were performed by the Center for Translational Genomics, Lund University, and Clinical Genomics Lund, SciLifeLab. Reads were aligned using STAR software (RRID:SCR_004463), and the reference genome sequence was from the Ensembl database, the human GRCh38 primary assembly, and the annotation (GTF) from GENCODE v33. Raw counts were normalized using DESeq2.

### Survival study: PCZ + PIT combination + COJEC

Injection of LU-NB-1 PDX cells (1 x 10^6^) was performed subcutaneously in NSG females as above. When the tumor reached 200-300 mm^3^, the mice were randomized into treatment groups: control, PCZ + PIT combination, COJEC, or PCZ + PIT combination + COJEC (**Fig. 5E**).

PCZ + PIT (total volume 50 µl; 5 mg/kg of each drug) was administered i.t. daily five times per week, as described for the i.t. study (**Fig. 2F**). The injection site was altered around the tumor to assure equal distribution across the tumor (**Fig. S6C**). All mice received i.t. injections on treatment days, i.e., vehicle (2.5% DMSO, 2.5% Tween 80, 40% PEG, 55% PBS) was injected in tumors in the control and COJEC-only groups. COJEC treatment was given i.p. and consisted of five different chemotherapies distributed in cycles over one week (**Fig. 5E**); Day 1 – cisplatin + vincristine, Day 3 – etoposide + cyclophosphamide, Day 5 – carboplatin (see **Fig. 5E** for doses). All COJEC drugs were dissolved in saline. Treatment breaks in the COJEC protocol were added if the mouse weight dropped below 90%.

Treatment was administered for 30 days, after which tumors were allowed to regrow. Mouse weights and tumor volumes were measured three times per week throughout the study. Mice were sacrificed when the tumors reached 1500 mm^3^. Tumors tissue was collected and snap frozen or fixed in 4% paraformaldehyde and embedded in paraffin.

### Statistical analysis

Data were analyzed in GraphPad Prism v9.3.0 for Windows (RRID:SCR_002798, GraphPad Software, San Diego, CA) or Excel 2016 (Microsoft, RRID:SCR_016137). *In vivo* groups were compared using one-way ANOVA on the last day of all mice being alive followed by Tukey’s multiple comparison test, unless stated otherwise. Survival analysis was performed with Kaplan-Meier analysis and log-rank tests between indicated groups. In cell state enrichment analysis, groups were compared with the Welch *t*-test. Mevalonate rescue experiments were analyzed using one-way ANOVA test and treatment groups were compared to control with Dunnet’s multiple comparison test. Live cell imaging experiment results were analyzed with one-way ANOVA followed by Tukey’s multiple comparison test between all groups.

## Acknowledgments

This work is dedicated to Neil T. Thompson, a great scientist and colleague who passed away during the finalization of this work. We thank the Center for Translational Genomics (CTG), Lund University and Clinical Genomics Lund, SciLifeLab for providing sequencing services. Part of the computations and data handling were enabled by resources provided by the National Academic Infrastructure for Supercomputing in Sweden (NAISS), partially funded by the Swedish Research Council through grant agreement no. 2022-06725. We thank Sebastian Braun from the Division of Translational Cancer Research, Lund University for sharing his confocal microscopy expertise. Figures 1, 3D, and 6 were created with BioRender.com.

## Funding

ENEA-European Neuroblastoma Association (DB),

aPODD Foundation (DB)

Healx (DB)

Swedish Cancer Society 20 0897 PjF and 23 2754 Pj (DB)

Swedish Childhood Cancer Foundation grants PR2020-0018 and PR2023-0006 (DB)

Swedish Research Council grants 2021-02597 and 2023-02402 (DB)

Crafoord Foundation grant 20210593 (DB)

Region Skåne and Skåne University Hospital Funding grant (DB)

Swedish Childhood Cancer Foundation grants TJ2021-0137 and PR2021-0129 (OCBR)

KI Forskningsbidrag grant 2022-01925 (OCBR)

Kungliga Fysiografiska Sällskapet i Lund 41984 and 42964 (KR)

## Author contributions

DBe, CS, and DBr conceptualized the study

DBe, DBr, KA, KR designed the study

DM, DO, IR, AL, JB, ED, DB, and NT performed the original predictions and drug selection at Healx.

KR, KA, AS, and KH performed *in vitro* experiments

KR, KA, JE, AA, AM performed *in vivo* experiments

KR and KA analyzed and visualized the data

EMO, DM, and OCBR performed bioinformatic analysis

OR and PS performed lipidomic experiments and analysis

KR, KA, and DB wrote the manuscript

DBe, CS, NT, DBr, and KA supervised the study

## Data and materials availability

Datasets created in this manuscript as well as it’s analyses will be made available at the time of publication (submission to array express in progress) and can be shared upon reviewer request.

