## Supplemental Figures and Tables for "Repurposing statins and phenothiazines to treat chemoresistant neuroblastoma"

Katarzyna Radke *et al.*

**This PDF file includes:**

Figs. S1 to S6
Tables S1 to S4

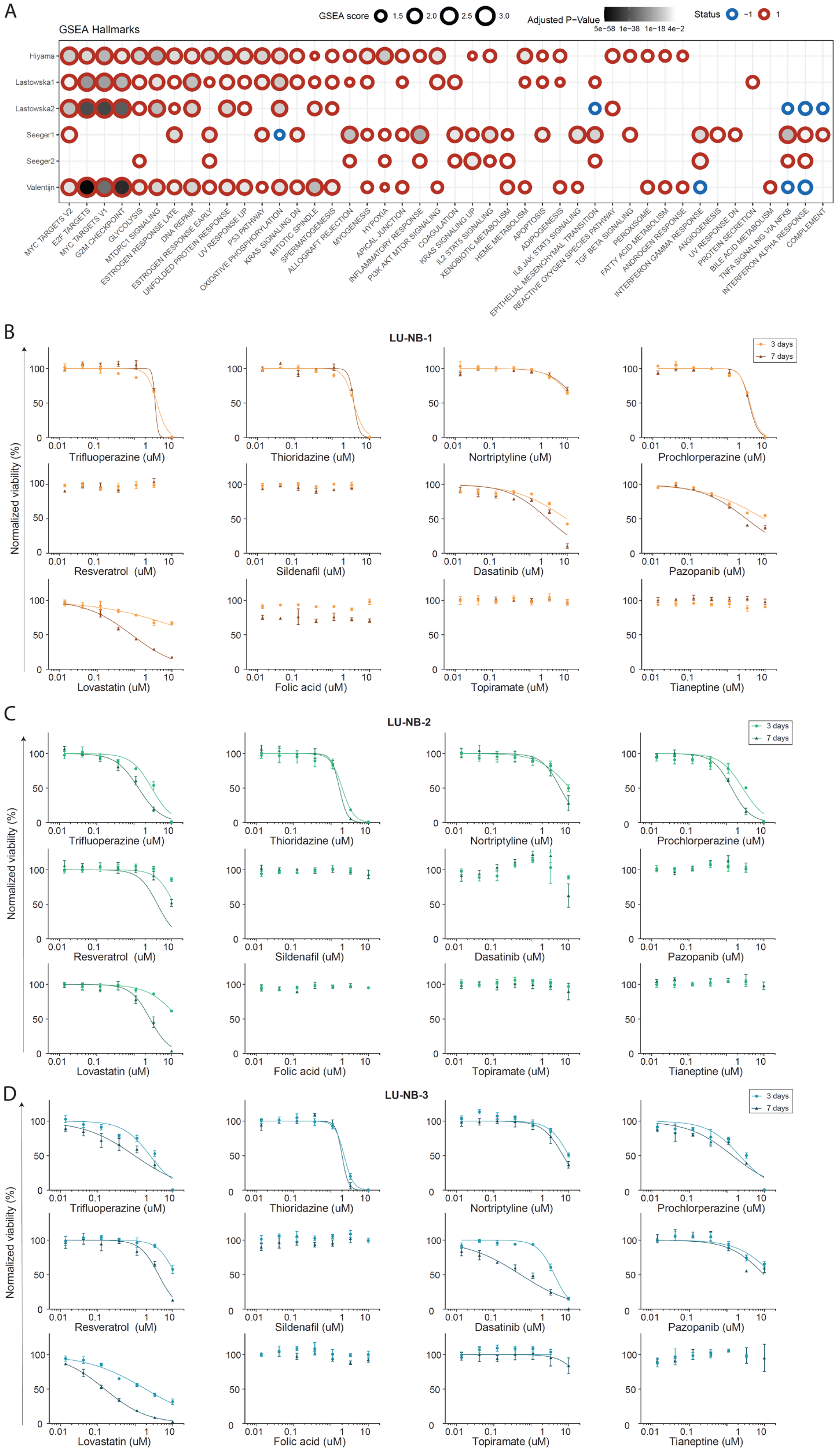
A
GSEA Hallmarks
GSEA score
1.5
2.0
2.5
3.0
Adjusted P−Value
5e−58
1e−38
1e−18
4e−2
Status
−1
1
Hiyama
Lastowska1
Lastowska2
Seeger1
Seeger2
Valentijn
MYC TARGETS V2
E2F TARGETS
MYC TARGETS V1
G2M CHECKPOINT
GLYCOLYSIS
MTORC1 SIGNALING
ESTROGEN RESPONSE LATE
DNA REPAIR
ESTROGEN RESPONSE EARLY
UNFOLDED PROTEIN RESPONSE
UV RESPONSE UP
OXIDATIVE PHOSPHORYLATION
P53 PATHWAY
KRAS SIGNALING DN
MITOTIC SPINDLE
SPERMATOGENESIS
ALLOGRAFT REJECTION
MYOGENESIS
HYPOXIA
APICAL JUNCTION
INFLAMMATORY RESPONSE
PI3K AKT MTOR SIGNALING
COAGULATION
KRAS SIGNALING UP
IL2 STAT5 SIGNALING
XENOBIOTIC METABOLISM
HEME METABOLISM
APOPTOSIS
ADIPOGENESIS
IL6 JAK STAT3 SIGNALING
EPITHELIAL MESENCHYMAL TRANSITION
REACTIVE OXYGEN SPECIES PATHWAY
TGF BETA SIGNALING
PEROXISOME
FATTY ACID METABOLISM
ANDROGEN RESPONSE
INTERFERON GAMMA RESPONSE
ANGIOGENESIS
UV RESPONSE DN
PROTEIN SECRETION
BILE ACID METABOLISM
TNFA SIGNALING VIA NFKB
INTERFERON ALPHA RESPONSE
COMPLEMENT
B
LU-NB-1
3 days
7 days
Normalized viability (%)
Trifluoperazine (uM)
Thioridazine (uM)
Nortriptyline (uM)
Prochlorperazine (uM)
Resveratrol (uM)
Sildenafil (uM)
Dasatinib (uM)
Pazopanib (uM)
Lovastatin (uM)
Folic acid (uM)
Topiramate (uM)
Tianeptine (uM)
C
LU-NB-2
3 days
7 days
Normalized viability (%)
Trifluoperazine (uM)
Thioridazine (uM)
Nortriptyline (uM)
Prochlorperazine (uM)
Resveratrol (uM)
Sildenafil (uM)
Dasatinib (uM)
Pazopanib (uM)
Lovastatin (uM)
Folic acid (uM)
Topiramate (uM)
Tianeptine (uM)
D
LU-NB-3
3 days
7 days
Normalized viability (%)
Trifluoperazine (uM)
Thioridazine (uM)
Nortriptyline (uM)
Prochlorperazine (uM)
Resveratrol (uM)
Sildenafil (uM)
Dasatinib (uM)
Pazopanib (uM)
Lovastatin (uM)
Folic acid (uM)
Topiramate (uM)
Tianeptine (uM)

20

**Fig. S1. Gene expression profile (GEP) analysis and *in vitro* response of predicted drugs.**

**A** Gene set enrichment analysis (Hallmarks database) of GEPs across datasets included in the original drug predictions (Table S1). **B-D** Single dose-response curves for the predicted 12 drugs (Table 1). NB PDX-derived organoids (n=3) were treated for 3 or 7 days: LU-NB-1, LU-NB-2, and LU-NB-3.

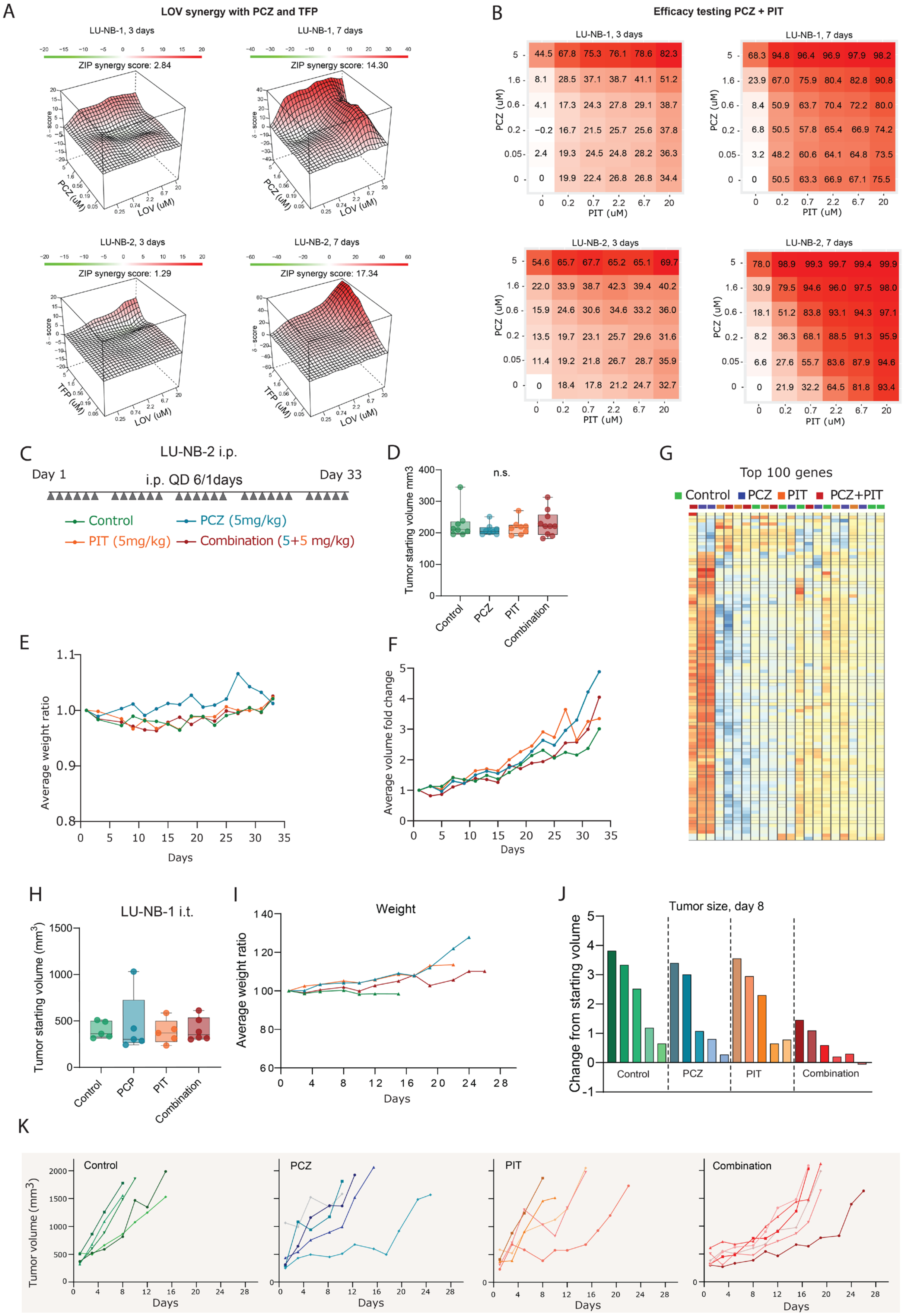
A
LOV synergy with PCZ and TFP
LU-NB-1, 3 days
ZIP synergy score: 2.84
LU-NB-1, 7 days
ZIP synergy score: 14.30
LU-NB-2, 3 days
ZIP synergy score: 1.29
LU-NB-2, 7 days
ZIP synergy score: 17.34
PCZ (uM)
TFP (uM)
LOV (uM)
B
Efficacy testing PCZ + PIT
LU-NB-1, 3 days
LU-NB-1, 7 days
LU-NB-2, 3 days
LU-NB-2, 7 days
PCZ (uM)
PIT (uM)
C
LU-NB-2 i.p.
Day 1
i.p. QD 6/1days
Day 33
Control
PCZ (5mg/kg)
PIT (5mg/kg)
Combination (5+5 mg/kg)
D
n.s.
Tumor starting volume mm3
Control
PCZ
PIT
Combination
E
Average weight ratio
Days
F
Average volume fold change
Days
G
Top 100 genes
Control
PCZ
PIT
PCZ+PIT
H
LU-NB-1 i.t.
Tumor starting volume (mm³)
Control
PCP
PIT
Combination
I
Weight
Average weight ratio
Days
J
Tumor size, day 8
Change from starting volume
Control
PCZ
PIT
Combination
K
Control
PCZ
PIT
Combination
Tumor volume (mm³)
Days

**Fig. S2**. **Evaluation of therapeutic efficacy of the chosen combination in NB**. **A** Heatmap representation (synergy matrix) based on cell viability after combination treatment with LOV and PCZ or TFP over 3 and 7 days (n=2). **B** Synergy matrix of the PCZ+PIT combination for LU-NB-1 and LU-NB-2 over 3 or 7 days (n=2). **C** Schematic overview of the LU-NB-2 treatment study *in vivo*. Drugs were administered intraperitoneally (i.p.) six times a week for 33 days. **D** Tumor starting volume (one-way ANOVA). **E** Average weight ratio. **F** Average tumor size in each treatment group throughout the study duration. **G** Unsupervised gene expression analysis of the top 100 most variable genes from tumor samples collected on day 33. **H-K** LU-NB-1 intra-tumoral (i.t.) study: **H** Tumor starting volume. **I** Average weight ratio. **J** Day 8 (last day of all mice alive): percentage of tumor size change from baseline for each mouse. **K** Tumor growth of individual mice over time.

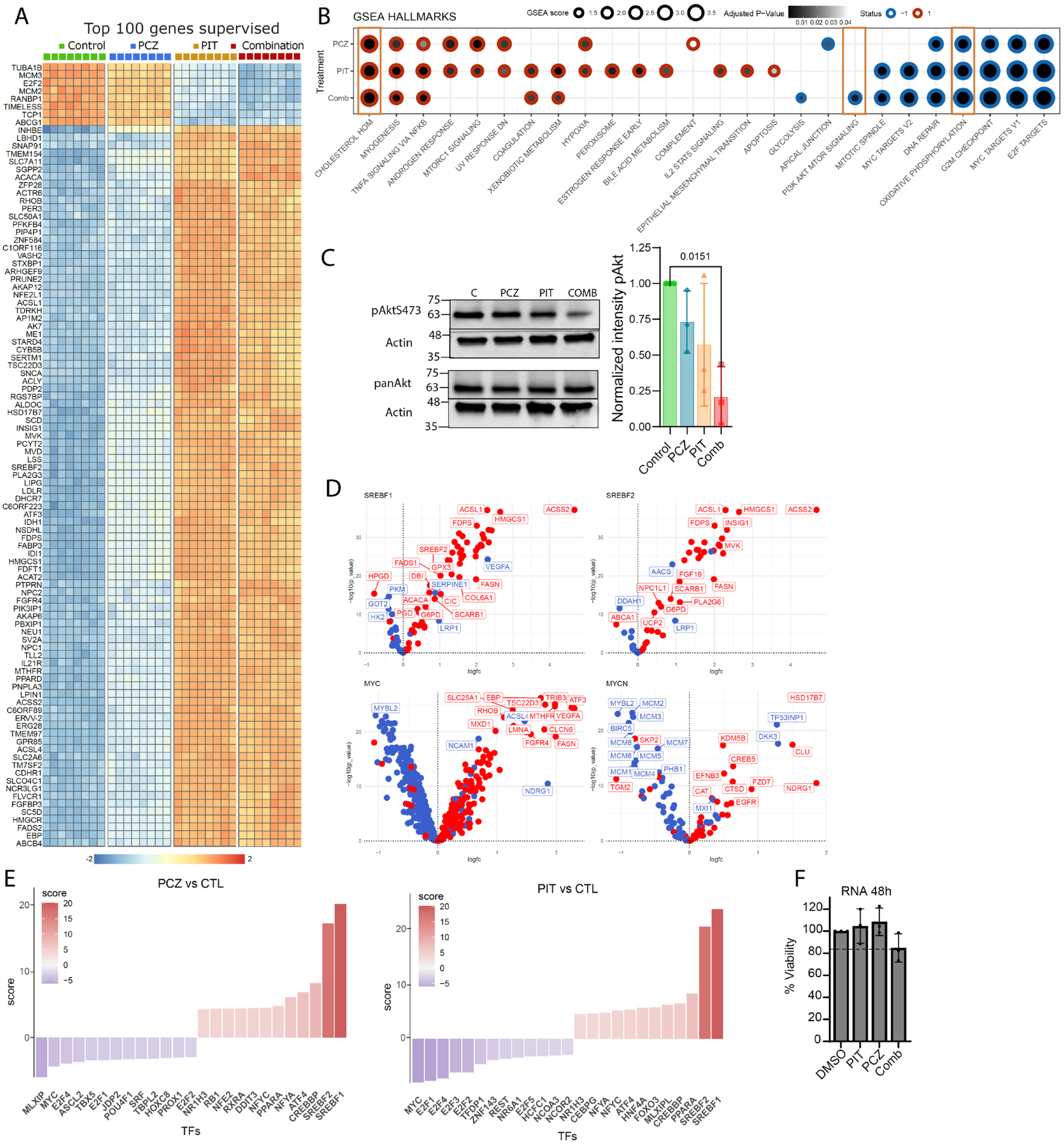


**Fig. S3. Mechanistic consequences of treatment with single drugs and PCZ + PIT combination.** RNA-seq and western blot analyses of LU-NB-2 organoids after 48 h treatment with single drugs or PCZ+PIT combination. **A** Heatmap showing differential gene expression between all treatment groups (top 100 most differentially expressed genes). **B** GSEA (Hallmarks database) of the respective treatment groups vs. control. **C** Western blot analysis of pAkt-S473 and panAkt protein expression (n=3, one-way ANOVA followed by Dunnet's multiple comparison). **D** Volcano plots displaying expression of the downstream targets of transcription factors *SREBF1*, *SREBF2*, *MYC*, and *MYCN* after PCZ+PIT combination treatment. **E** Transcription factor (TF) activity after treatment with single drugs (CollecTri). **F** Viability of LU-NB-2 PDX organoids at collection for RNA-seq after 48 h treatment with the respective drugs and with the combination (n=3, one-way ANOVA, p=0.154).

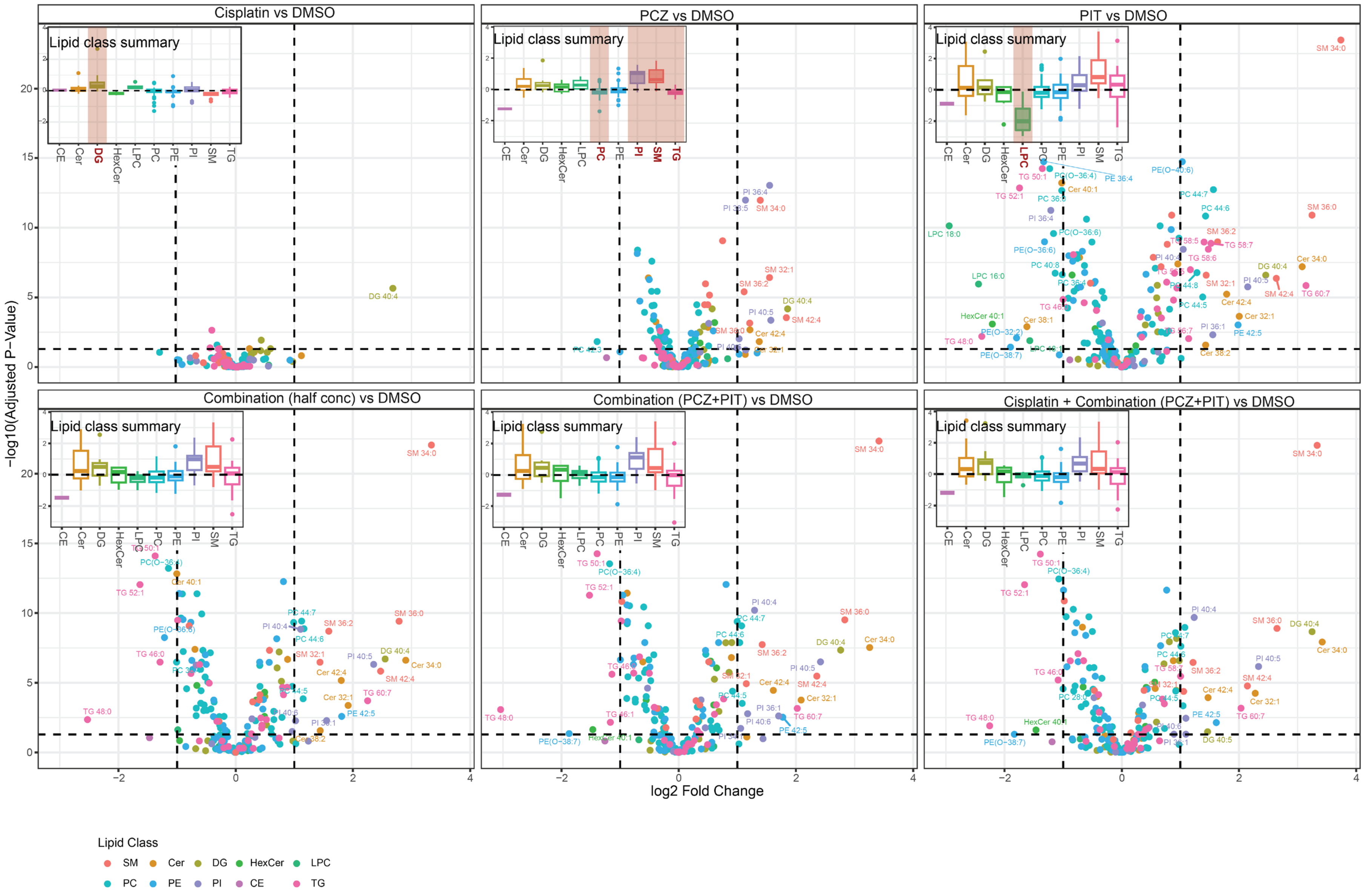

Cisplatin vs DMSO
PCZ vs DMSO
PIT vs DMSO
Combination (half conc) vs DMSO
Combination (PCZ+PIT) vs DMSO
Cisplatin + Combination (PCZ+PIT) vs DMSO
Lipid class summary
−log10(Adjusted P−Value)
log2 Fold Change
Lipid Class
SM
Cer
DG
HexCer
LPC
PC
PE
PI
CE
TG

**Fig. S4. Lipidomics changes after single drugs, combination, and combination + cisplatin treatment *in vitro.*** Volcano plot presenting upregulated and downregulated lipids (species level nomenclature) after treatment with cisplatin, PCZ, PIT, PCZ+PIT at halved and full concentrations and cisplatin+PCZ+PIT. Upper left corner of each panel shows logFC summary of each lipid class change after treatment and significant changes were indicated with the red background. DMSO treated cells were used as a baseline.

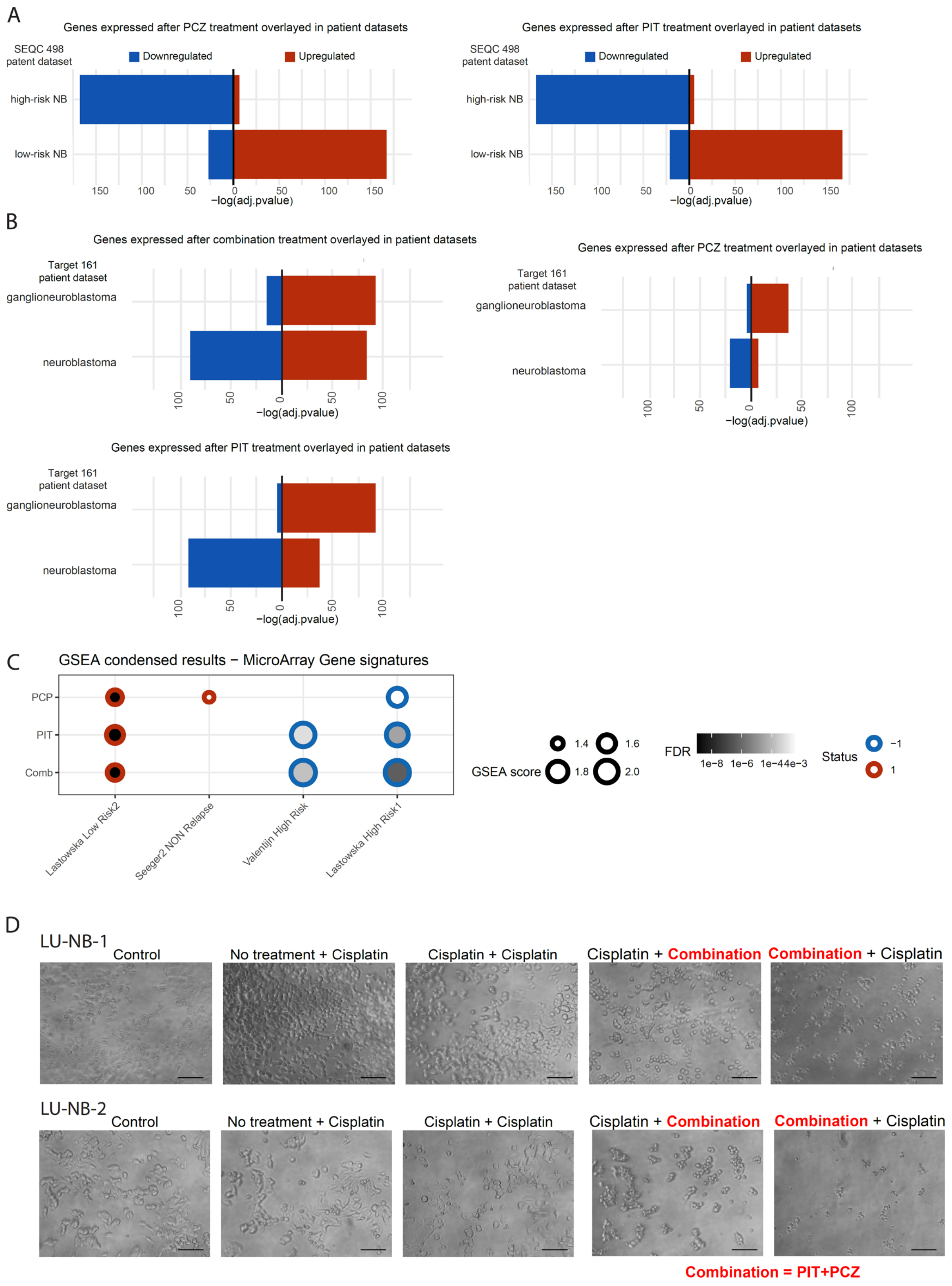
A
Genes expressed after PCZ treatment overlayed in patient datasets
SEQC 498
patent dataset
Downregulated
Upregulated
high-risk NB
low-risk NB
−log(adj.pvalue)
Genes expressed after PIT treatment overlayed in patient datasets
B
Genes expressed after combination treatment overlayed in patient datasets
Target 161
patient dataset
ganglioneuroblastoma
neuroblastoma
Genes expressed after PCZ treatment overlayed in patient datasets
Genes expressed after PIT treatment overlayed in patient datasets
C
GSEA condensed results − MicroArray Gene signatures
PCP
PIT
Comb
Lastowska Low Risk2
Seeger2 NON Relapse
Valentijn High Risk
Lastowska High Risk1
GSEA score
FDR
Status
D
LU-NB-1
Control
No treatment + Cisplatin
Cisplatin + Cisplatin
Cisplatin + Combination
Combination + Cisplatin
LU-NB-2
Combination = PIT+PCZ

**Fig. S5. Transcriptional phenotype correlated with risk in patient datasets. A** Association of DEGs (after single PCZ and PIT treatment) with risk stratification in the SEQC498 patient dataset. **B** Genes expressed after PCZ and/or PIT treatment overlayed in the Target161 dataset including patients with ganglioneuroblastoma and patients with aggressive neuroblastoma. **C** GSEA of RNA expression after treatment with single drugs and combination (LU-NB-2, 48 h) compared with risk in original datasets (Table S1). Only significant (FDR p-value $< 0.05$) results are displayed. **D** Brightfield images of cells seeded on laminin at different treatment conditions. Corresponding pictures of normalized viability graphs in Fig 4E. Picture magnification: 10x, scale bar 100 µm.

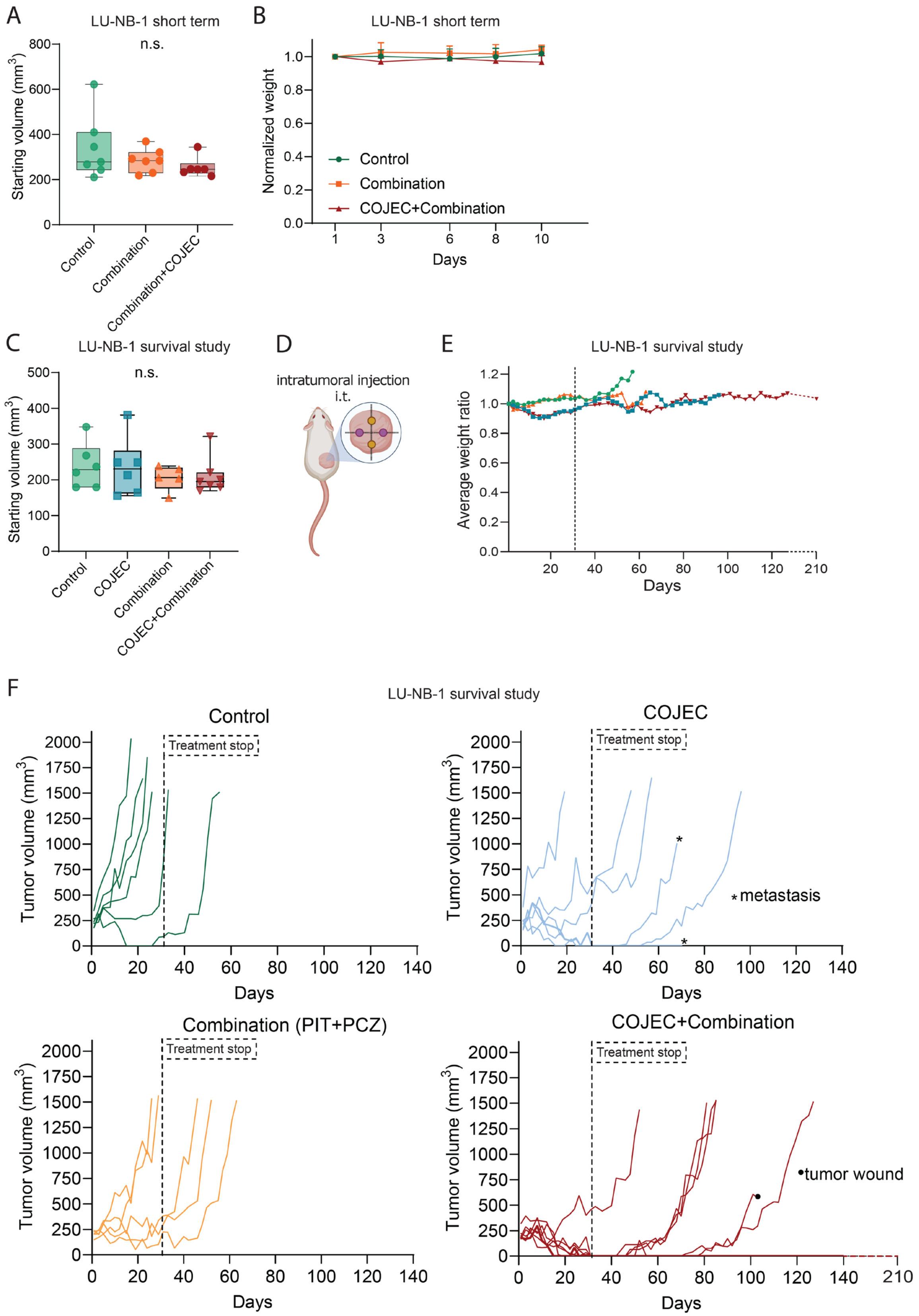
A
LU-NB-1 short term
n.s.
Starting volume (mm³)
Control
Combination
Combination+COJEC
B
LU-NB-1 short term
Normalized weight
Control
Combination
COJEC+Combination
Days
C
LU-NB-1 survival study
n.s.
Starting volume (mm³)
Control
COJEC
Combination
COJEC+Combination
D
intratumoral injection
i.t.
E
LU-NB-1 survival study
Average weight ratio
Days
F
LU-NB-1 survival study
Control
Treatment stop
Tumor volume (mm³)
Days
COJEC
Treatment stop
* metastasis
Combination (PIT+PCZ)
Treatment stop
COJEC+Combination
Treatment stop
• tumor wound

**Fig. S6. *In vivo* studies of PCZ+PIT drug combination together with COJEC chemotherapy. (A-B)** Short-term study (10-day treatment), LU-NB-1 model: **A** Starting tumor volume in all groups. **B** Mouse weight during treatment. **(C-F)** Survival study (30 d treatment) in LU-NB-1 tumors: **C** Starting tumor volume in all groups. **D** Schematic of i.t. injection approach. **E** Average weight throughout the study. **F** Single mouse tumor size per group, over time.

**Table S1. Neuroblastoma expression datasets used for drug predictions.**

| **Author** | **NCBI GEO** | **Reference name** | **Benign group** | **Progression/higher severity group** | **Patient group** | **Number of patients** |
|---|---|---|---|---|---|---|
| Valentijn et al. (2015) *(87)* | GSE73537 | Valentijn | Alive | Dead | All stages | 34 |
| Ohtaki et al. (2010)*(88)* | GSE16237 | Hiyama | Alive + stages 1/2/4s | Dead of disease + stages 3/4 | All stages | 51 |
| Asgarzadeh et al.(2006) *(89)* | GSE3446 HG-U133A | Seeger 1 | No relapse | Relapse/metastasis | *MYCN* non-amplified | 117 |
| Asgarzadeh et al.(2006) *(89)* | GSE3446 HG-U133B | Seeger 2 | No relapse | Relapse/metastasis | *MYCN* non-amplified | 117 |
| Lastowska et al.(2007)*(90)* | GSE13136 | Lastowska 1 | Low stage, MYCN non-amplified | *MYCN* amplified, high stage, 1p del | All stages, 17q gain | 30 |
| Lastowska et al. (2007) *(90)* | GSE13136 | Lastowska 2 | Low stage, *MYCN* non-amplified | *MYCN* non-amplified, high stage, 11q del | All stages, 17q gain | 30 |

**Table S2. Neuroblastoma PDX models included in the study.**

| PDX | Sample type | SNP profile | Stage | Chemotherapy |
|---|---|---|---|---|
| LU-NB-1 | Primary (adrenal gland) | *MYCN* amp, 1p del, 17q gain | IV | No |
| LU-NB-2 | Metastasis (after relapse) | *MYCN* amp, 1p del, 17q gain | IV | Yes |
| LU-NB-3 | Primary (adrenal gland) | *MYCN* amp, 1p del, 17q gain | III | No |

**Table S3**. **Effect of four selected drugs in three different high-risk neuroblastoma organoid models.** Single dose curves for statistical comparison between 3 and 7 days. NA- not enough data points around $IC_{50}$ to compute the CI.

| | LU-NB-1 | | LU-NB-2 | | LU-NB-3 | |
|---|---|---|---|---|---|---|
| | 3 days | 7 days | 3 days | 7 days | 3 days | 7 days |
| **Trifluoperazine** | | | | | | |
| $IC_{50}$ | 4.01 | 3.57 | 2.95 | 1.31 | 2.50 | 0.96 |
| 95% CI $IC_{50}$ | NA | NA | 2.4 – 3.6 | 1.1 – 1.5 | 1.8 – 3.4 | 0.62 – 1.49 |
| Sig 3 vs 7 days | no | no | yes | yes | yes | yes |
| AUC | 495 | 542 | 427 | 240 | 408 | 304 |
| 95% CI AUC | 486 – 504 | 521 – 563 | 390 – 63.4 | 211 – 268 | 376 – 441 | 265 – 342 |
| Sig 3 vs 7 days | yes | yes | yes | yes | yes | yes |
| **Thioridazine** | | | | | | |
| $IC_{50}$ | 3.78 | 3.81 | 1.99 | 1.67 | 2.25 | 1.99 |
| 95% CI $IC_{50}$ | 3.53 – 4.06 | NA | 1.70 – 2.33 | NA | 2.06 – 2.50 | NA |
| Sig 3 vs 7 days | no | no | no | no | no | no |
| AUC | 478 | 533 | 278 | 224 | 300 | 247 |
| 95% CI AUC | 468 – 487 | 521 – 545 | 258 – 298 | 205 – 244 | 270 – 331 | 212 – 283 |
| Sig 3 vs 7 days | yes | yes | yes | yes | no | no |
| **Prochlorperazine** | | | | | | |
| $IC_{50}$ | 3.93 | 3.77 | 2.73 | 1.38 | 2.20 | 1.41 |

| | | | | | | |
|---|---|---|---|---|---|---|
| 95% CI $IC_{50}$ | NA | NA | 2.08 – 3.50 | 1.24 – 1.53 | 1.48 – 3.16 | 0.87 – 2.23 |
| Sig 3 and 7 days | no | no | yes | yes | no | no |
| AUC | 498.2 | 488.2 | 409.6 | 232.4 | 393.0 | 333.4 |
| 95% CI AUC | 486 – 510 | 477 – 499 | 388 – 432 | 199 – 266 | 363 – 423 | 318 – 349 |
| Sig 3 vs 7 days | no | no | no | no | no | no |
| **Lovastatin** | | | | | | |
| $IC_{50}$ | 45.9 | 0.84 | 15.9 | 2.67 | 1.89 | 0.15 |
| 95% CI $IC_{50}$ | 24.0 – 119 | 0.73 – 0.97 | 13.0 – 21.4 | 2.30 – 3.09 | 1.54 – 2.33 | 0.14 – 0.17 |
| Sig 3 vs 7 days | yes | yes | yes | yes | yes | yes |
| AUC | 700 | 303 | 796 | 398 | 427 | 106 |
| 95% CI AUC | 670 – 730 | 290 – 315 | 780 – 811 | 343 – 453 | 396 – 458 | 97.2 – 115 |
| Sig 3 vs 7 days | yes | yes | yes | yes | yes | yes |

**Table S4. Neuroblastoma gene signatures included in the analysis.**

| Author | Material | States | Omics method |
|---|---|---|---|
| Van Groningen *et al.* 2017*(12)* | Cell lines (N=30)<br>Adr-mes cell line pairs (N=4) | Adrenergic<br>Mesenchymal | Chip-Seq, RNA-seq, |
| Boeva *et al.* 2017*(13)* | Patient tumors (N=10)<br>NB cell lines (N=25)<br>PDXs (N=6) | Noradrenergic<br>NCC-like | Chip-Seq, RNA-seq, |
| Gartlgruber *et al.* 2021*(16)* | Patient tumors (N=60)<br>Cell lines (N=25) | *MYCN* amp<br>*MYCN* non-amp, high risk<br>*MYCN* non-amp, low risk<br>MES/Schwann cell precursor | Chip-Seq (H3K27ac), RNAseq, ATAC-seq |
| Olsen *et al.* bioRxiv*(18)* | Patient tumors (N=17) | Adrenergic<br>Mesenchymal<br>SCP-like<br>bridge cells | scRNA-seq |
| Manas *et al.* 2022*(19)* | Integration of signatures from PDX models (N=3) and multiple published signatures | Adrenergic<br>Mesenchymal-like | RNA-seq |
| Bedoya-Reina *et al.* 2021*(15)* | Patient tumors (N=11)<br>Normal adrenal gland (human+mouse) (N=3+5) | Undiff (nC2,3)<br>NOR (nC5,7,8,9)<br>Stromal clusters | sc/snRNA-seq |
| Yuan *et al.* 2022*(17)* | Patient tumors (N=10 peripheral neuroblastic tumors) | Adrenergic,<br>Transitional<br>Mesenchymal | scRNA-seq |
| Patel *et al.* bioRxiv*(22)* | Patient tumors (N=51)<br>PDX1(N=1) | Adrenergic<br>Mesenchymal<br>Sympathoblast | sc/snRNA-seq<br>Spatial transcriptomics |
